# Parent-of-origin effect on global gene expression and host-plant adaptation in *Spodoptera frugiperda* (Lepidoptera: Noctuidae)

**DOI:** 10.1101/2025.03.10.642409

**Authors:** Laijiao Lan, Stéphanie Robin, Kiwoong Nam, Nicolas Nègre

**Affiliations:** DGIMI, Univ Montpellier, INRAE, Montpellier, France; BIPAA, IGEPP, INRAE, Institut Agro, University of Rennes, Rennes, France; IRISA, University of Rennes, INRIA, CNRS, IRISA, Rennes, France

**Keywords:** *Spodoptera frugiperda*, RNA-seq, maternal effect, heterosis effect, oxidative phosphorylation

## Abstract

*Spodoptera frugiperda* (Fall armyworm, FAW), is a major pest that adapt to many host plants and cause significant economic losses. There are two genetically differentiated FAW strains with different food preferences, the “corn strain” (SfC) and the “rice strain” (SfR). Large transcriptional differences between the strains have been previously documented, which suggested a shift in metabolic homeostasis. In this study, we investigated how parental inheritance can influence FAW metabolism by comparing the larval performance as well as gene expression between FAW strains and their reciprocal hybrids. We recorded life history traits of these genotypes, including larval weight and larval survival rate, when reared on artificial diet or on corn leaves. We observed a previously documented heterosis effect with hybrids gaining more weight than parental lines on both diets. With corn diet, we also observed a significant maternal influence on fitness between the hybrids. We then performed whole genome sequencing and RNA-sequencing on individuals from the same artificial rearing conditions to correlate these differences in fitness with variations on gene expression. We found that the heterosis effect is potentially associated with differentially expressed genes predominantly engaged in immune response and metabolism. The two different hybrid genotypes differed in the expression of RNA gene silencing and oxidative phosphorylation, suggesting a strong effect of parent-of-origin variations on global metabolic homeostasis, as well as a potential involvement of transposable element silencing in compatibilities between the FAW strains genomes, which might explain the differences in FAW plant adaptation.

## Introduction

Environmentally induced variations in gene expression can act as a fundamental mechanism of phenotypic plasticity (Schlichting and Smith, 2002), which enables organisms to adapt rapidly to environmental changes (West-Eberhard, 1989; Fordyce, 2006; Pigliucci *et al*., 2006). It can also enable generalist herbivores to contend with a broader range of food sources (Schneider *et al*., 2024; Sharma *et al*., 2024). In contrast, the presence of more specialized biotypes among phytophagous insects provides valuable systems for investigating the genomic basis of host-plant adaptation (Drès and Mallet, 2002; Forister *et al*., 2012; Eyres *et al*., 2016). While genetic variations may be the basis for adaptative evolution to plants, their impact on global gene expressions and thus on plasticity is harder to disentangle.

*Spodoptera frugiperda* (J. E. Smith) (Lepidoptera: Noctuidae), also known as the fall armyworm (FAW), is a major worldwide agricultural pest (Kenis *et al*., 2023). FAW is highly polyphagous, infesting a range of over 350 plant species across 76 distinct botanical families, with a particular preference for graminaceous staple crops (Montezano *et al*., 2018; Malo and Hore, 2020), such as corn and sorghum. Despite its polyphagy, FAW is also known to be constituted of two strains: the corn strain (SfC) and the rice strain (SfR) (Pashley *et al*., 1985). While morphologically identical, the two FAW strains have different host-plant preferences (Pashley, 1986, 1988) and show significant differences in growth and reproductive behaviors (Schöfl *et al*., 2009; Orsucci *et al*., 2022; Lan and Nègre, 2023). Several genetic markers have been utilized to identify the two strains, among which the maternally inherited mitochondrial *COI* gene (Cytochrome c Oxidase subunit I) (Levy *et al*., 2002; Nagoshi *et al*., 2006) and the Z-chromosome-located *Tpi* gene (Triose-phosphate isomerase) (Nagoshi, 2010) being the most applied. Phylogenetic analyses employing mitochondrial DNA markers have delineated distinct clades corresponding to the two strains (Kergoat *et al*., 2012; Dumas, *et al*., 2015; Gouin *et al*., 2017), and genome-wide analyses show that the divergence between them is indicative of incipient speciation (Dumas *et al*. 2015; Gouin *et al*., 2017; Fiteni *et al*., 2022). Indeed, laboratory studies have successfully documented instances of mating between the two distinct strains of FAW (Velásquez-Vélez *et al*., 2011; Dumas, *et al*., 2015; Lan and Nègre, 2023). Moreover, field populations have been found to contain a high prevalence of hybrids (Prowell *et al*., 2004). Whole-genome resequencing data and genome-wide SNP (single nucleotide polymorphisms) analyses showed whole-genome differentiation but only a handful of loci targeted by selective sweeps between the FAW strains (Fiteni *et al*., 2022). Nonetheless, extensive transcriptional differences have been identified between the two strains of FAW with laboratory as well as field populations (Roy *et al*., 2016; Orsucci *et al*., 2022). Intriguingly, our group identified a pronounced and consistent difference between strains at the level of mitochondrial transcription, which may reflect fundamental metabolic and physiological distinctions between the strains (Orsucci *et al*., 2022), resulting in their different adaptation to host plants. However, the specific molecular mechanisms affected by this variation in mitochondrial expression are not yet understood.

In this study, we investigate the potential influence of maternally inherited loci (particularly the mitochondrial genes) on the larval performance of FAW and global transcriptional patterns, thereby affecting their adaptability to different host plants. We established four distinct FAW genotypes: two parental strains (CC: SfC♀ × SfC♂, RR: SfR♀ × SfR♂), and their reciprocal hybrids (CR: SfC♀ × SfR♂, RC: SfR♀ × SfC♂). Ideally, if the parental strains were isogenic, the hybrids should have identical heterozygous genotypes for all genetic loci except in their mitochondrial DNA and sex chromosomes due to different parent of origins. We reared the four populations simultaneously on artificial diet and corn plants under individually rearing condition and observed a strong heterosis effect, with hybrids outperforming parental lines in weight gain, consistent with previous findings from mass rearing (Lan and Nègre, 2023). Moreover, a significant maternal influence on fitness between the hybrids was also observed on corn diet. We then combined these performance experiments with DNA-seq and RNA-seq to analyze the transcriptional plasticity explaining the variations in life history traits. Indeed, we observed a strong impact of genotypes on global transcriptional patterns. The observed heterosis effect potentially involves a cross-talk between immunity and metabolism in hybrids. The differences between reciprocal hybrids showed the involvement of RNAi machinery in the hybrids, suggesting genomic incompatibilities between them. These global changes in transcription between genotypes are likely due to overall homeostatic imbalance compensation.

## Materials and Methods

### Biological materials

Two laboratory FAW host strains were used: corn strain (SfC) and rice strain (SfR). The SfC population was seeded with individuals collected from Guadeloupe in 2001. The SfR population was seeded with 50 pupae collected in Florida (Hardee County) in 2012. Both strains have been reared under laboratory conditions for many generations, at a temperature of 24 ± 1°C (T) with a 16: 8h light: dark photoperiod (L: D) and 70% relative humidity (RH), on an artificial diet called Poitout (Poitout *et al*., 1972). The genomes of these two populations have been sequenced (Gouin *et al*., 2017) and are available on a dedicated database (https://bipaa.genouest.org/sp/spodoptera_frugiperda_corn/). FAW is considered as a quarantine pest in metropolitan France, thus all experiments below were conducted under a confined environment on an insect quarantine platform (PIQ, University of Montpellier, DGIMI laboratory, https://dgimi.hub.inrae.fr/plateforme-experimentale-sur-insectes).

### Experimentation

#### Performance experiment with Poitout/corn diet

Following biological containment procedures, we first performed reciprocal crosses between the two strains to establish two hybrid genotypes, SfR ♀ × SfC ♂ (RC), and SfC ♀ × SfR ♂ (CR). In the meantime, we maintained the two parental strains, SfR ♀ × SfR ♂ (RR), and SfC ♀ × SfC ♂ (CC). The four genotypes were produced in triplicate for a total of 12 populations. Each larva was reared individually on an artificial diet (Poitout) or on segments of fresh corn leaves in a 30ml transparent plastic vessel (4.4 × 4.4 × 3.2 cm) to avoid cannibalism. Each set of 30 vessels constituted a single replicate, organized within a tray for experimental uniformity. Ventilation was achieved by perforating the vessel lids with three small apertures. To prevent contamination and larval escape, each vessel was enclosed with a fine insect-proof mesh (150μm sieve) and a layer of sterile tissue. The replicate trays, three per genotype, were positioned at random on three distinct shelves within a controlled environmental incubator in PIQ (T = 25℃, RH = 40 ± 5%, L: D = 16: 8) to avoid the floor and edge effects. Upon reaching the end of L6 larval stage, characterized by feeding cessation and minimal locomotion, larvae were meticulously transferred with a soft tweezer to another sterilized 30ml plastic vessel containing a substrate of one-third vermiculite to stimulate pupal metamorphosis.

Throughout the larval development, we meticulously recorded several life history traits for each of the four genotypes, including temporal changes in larval weight and survival rates. Larval weight was quantified by individually weighing 10 randomly selected larvae from each replicate, totaling 30 larvae per genotype, at intervals from the 3rd instar stage through to pupation. Larval instar progression was ascertained by measuring the head capsule width, in accordance with established methodologies (Orsucci *et al*., 2022). Survival rates were calculated by comparing the number of larvae that survived to the end of the larval stage, just prior to metamorphosis, with the initial population size at the onset of the experiment. Larval survival rate was measured from the 1st larval instar to metamorphosis for the four genotypes, which was the number of surviving larvae at the end of the larval stage (before metamorphosis) that was counted over the initial number of larvae.

#### Reproduction comparison experiment

To determine the reproductive capacity of the four FAW genotypes, we conducted mating experiments. We compared the average offspring of two parental genotypes (RR ♀ × RR ♂, CC ♀ × CC ♂) with those of two F_1_ hybrid populations (RC ♀ × RC ♂, CR ♀ × CR ♂). For each experimental genotype, we placed five adult females and five males, which emerged simultaneously, in a 2L oblong plastic container to enhance mating opportunities. We then calculated the average number of offspring produced per female for each genotype to compare. The containers were ventilated with approximately 50 small holes in the lids. To prevent cross-contamination between populations and to keep the tiny neonate offspring from escaping, we covered the containers with an insect-proof mesh (150μm sieve), an additional layer of paper, and secured them. Inside each container, we provided a small plastic vial containing a 10% honey solution on a wet cotton pad for nutrition and a long strip of filter paper around the vial to stimulate oviposition. All experimental setups were maintained under identical conditions on the same shelf within an incubator (T = 25℃, HR = 40 ± 5%, L: D = 16: 8). The number of viable offspring from each setup was counted and analyzed to assess any differences in reproductive performance.

#### DNA and RNA extraction

Genomic DNA and total RNA were isolated from individual FAW larva of the four genotypes with AllPrep DNA/RNA Mini Kit (50) (Qiagen Cat. ID: 80204) in accordance with the manufacturer’s protocol. The DNA and RNA extraction process commenced by placing a single 4th instar larva into a 2 ml Eppendorf tube pre-filled with Buffer RLT Plus. This procedure was replicated for 3 larvae from each of the four distinct genotypes—CC, RR, CR, and RC—yielding a total of 12 samples (3 samples per genotype multiplied by 4 genotypes). Each larva was then subjected to mechanical disruption and homogenization using the TissueLyser II from Qiagen (Catalog ID: 85300), with the aid of a 5 mm bead to enforce thorough mixing within the tube. Post-homogenization, the lysate was carefully transferred to an AllPrep DNA spin column that had been positioned in a 2 ml collection tube and centrifuged to initiate the DNA purification process. The DNA-containing solution within the AllPrep DNA spin column was reserved for future DNA-related applications. The flow-through from this initial DNA purification step was directed towards a RNeasy spin column, which was utilized for the purification of total RNA. This dual approach allowed for the concurrent extraction of both DNA and RNA from the same starting material, streamlining the process and maximizing the use of each sample. Furthermore, to ensure that the RNA sample was free from any potential DNA contamination, we employed the TURBO DNA-*free*™ Kit (Invitrogen, Cat. ID: AM1907). This kit effectively removes contaminating DNA and eliminates DNase enzymes and divalent cations from the RNA sample, thus providing clean RNA preparation. After the purification process, the integrity and purity of both the DNA and RNA samples were evaluated using a UV spectrophotometer, specifically the NanoDrop device (Wilmington, DE, USA). The purity of nucleic acids was determined by measuring the absorbance ratios at 260 nm and 280 nm (A260/A280). For DNA, a ratio of 1.0 corresponds to a concentration of 50 µg/ml, indicating pure DNA, while for RNA, a ratio of 1.0 corresponds to a concentration of 40 µg/ml, which is indicative of pure RNA. These measurements ensure that the extracted nucleic acids are of high quality and suitable for downstream applications.

### Whole genome sequencing pipeline

DNA samples were sent for whole genome sequencing (DNA sequencing, DNA-seq). Paired-end libraries and 2*150 bp Illumina sequencing were performed by Novogene Europe (Cambridge, UK) on the NovaSeq S6000 platform. Fastq files were then aligned onto the SfC reference genome v7 (https://bipaa.genouest.org/sp/spodoptera_frugiperda_corn/) – also available on NCBI (https://www.ncbi.nlm.nih.gov/datasets/genome/GCA_019297735.2/) (Yainna *et al*., 2022), using bowtie2 v2.3.4.1 (Langmead and Salzberg, 2012), with the default parameters except the --very-sensitive-local parameter. Duplicate reads were then removed with Picard tools (https://broadinstitute.github.io/picard/) MarkDuplicates command.

We called genetic variations using haplotype calling with GATK v4.1.2.074 (McKenna *et al*., 2010). Only single nucleotide variations (SNVs) were retained, and filtering was applied to exclude SNVs meeting any of the following criteria: QD < 2.0, FS > 60.0, MQ < 40.0, MQRankSum < −12.5, or ReadPosRankSum < −8.0. If the genotype was not determined in at least 80% of the samples at a position, that position was also discarded.

### RNA-seq pipeline

Library construction and high throughput sequencing were performed for the RNA samples using Illumina technologies by Novogene Europe (Cambridge, UK) on a HiSeq 2000 (Illumina). The transcriptome reads were subjected to nf-core/rnaseq bioinformatics workflow (v3.10.1, https://nf-co.re/rnaseq, Ewels *et al*., 2020) for quality control, mapping and counting with default parameters. The mapping was done with STAR (v2.7.9a, Dobin *et al*., 2013), with between 73.69% and 86.65% of reads mapped only once. The counting was done with featureCounts with the default parameters except the following parameters: -g gene_id -t exon -C -p -s 0 -M --fraction (v2.0.1, Liao *et al*., 2014), with 65.1% to 77.3% assignment. The RNA-seq data of all the samples was aligned against a common reference, the *Spodoptera frugiperda* corn assembly v7.0 (Fiteni *et al*., 2022), and the counts were generated from a common structural annotation, the *Spodoptera frugiperda* corn annotation (OGS7.1_20230306) on BIPAA (https://bipaa.genouest.org/sp/spodoptera_frugiperda_corn/). The specific RNA-seq analysis workflow can be seen in **Supplementary Method** and **Supplementary Figure S14**.

### Statistical analysis

Statistical tests were carried out with R software (version 4.3.2, R Core Team, 2023), followed by residual analyses to verify the suitability of the distributions of the datasets. The larval weight and larval survival rate were analyzed using one-way analysis of variance, with FAW genotype (4) as the factor. The means were separated and compared with Tukey’s HSD test. The differential expression analysis was conducted with DESeq2 (v1.42.0), and the enrichment analysis was carried out with clusterProfiler (v4.10.0). Multiple other R packages were used (tidyverse, RColorBrewer, ggpubr, grid, DEGreport, gson, enrichplot, DOSE, ggplot2, cols4all, patchwork, dplyr, multcomp, multcompView, ggthemes, gplots, apeglm, etc.). All the codes and analysis files are publicly available on the accompanying **Supplementary Material** on Zenodo: 10.5281/zenodo.14856499.

## Results

### Performance differences among the four FAW genotypes

#### Larval growth and survival differences

When reared on artificial diet (Poitout), the two FAW hybrid populations CR and RC outperformed the two parental strains in weight throughout the larval stage (**Fig. 1A**). On both diets, CR was systematically the heaviest among all the genotypes for all the measurements (**Fig. 1A-B**).

**Figure 1.**
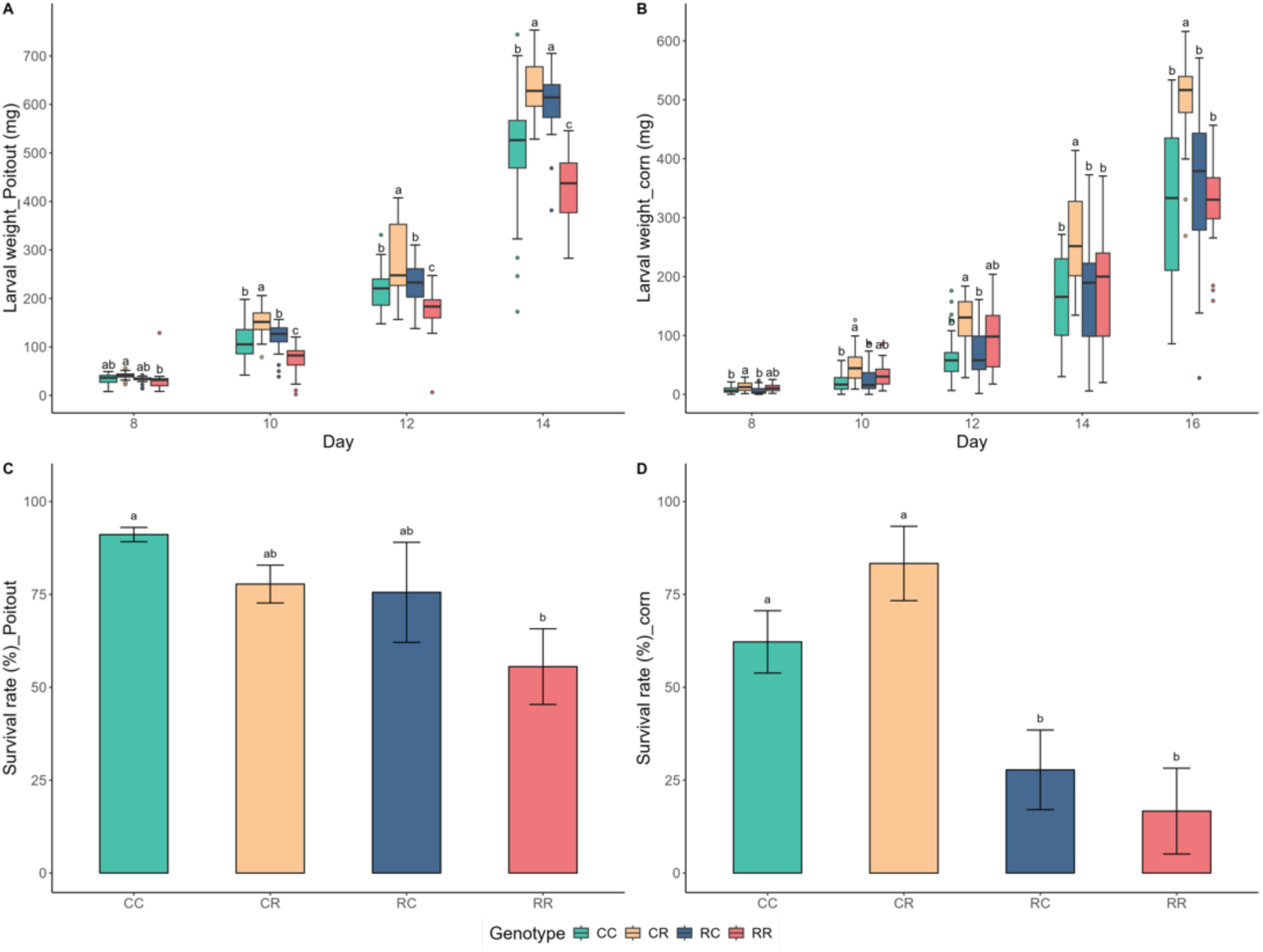
Larval performance differences among four FAW genotypes with artificial diet and corn diet. (A) Larval weight across the developmental time on artificial diet (Poitout); (B) Larval weight across the developmental time on corn diet; (C) Mean larval survival rate ± SD on Poitout; (D) Mean larval survival rate ± SD on corn diet. Larval weight measurements were done every other day from 3rd instar larvae until pupal metamorphosis (d8, d10, d12, and d14 on Poitout, d8, d10, d12, d14, and d16 on corn diet). The number of larvae recorded from the three replicates for each genotype was about (n = 90) in total. Different letters above bars indicate significant differences among these genotypes (Tukey’s HSD test, *p* < 0.05).

As for the larval survival rate, the two hybrid genotypes, CR and RC, had slightly lower survival rates than CC and higher survival rates than RR when reared on Poitout, though the differences were not significant (*p* > 0.05). On corn diet, CR had the highest survival rate and showed no significant difference from CC, while both RC and RR had significantly low survival rates compared to CC and CR (*p* < 0.05). These results confirm the heterosis effect observed previously on mass rearing conditions on artificial diet (Lan and Nègre, 2023) as well as the better performance of SfC mitotypes on corn (Orsucci *et al*., 2022).

#### Reproduction differences

A comparative inbreeding analysis of average progeny counts between the four populations was conducted (**Supplementary Fig. S1**). When reared on Poitout, all four genotypes successfully produced viable offspring, with the hybrid populations crosses (RC × RC and CR × CR) yielding a notably elevated mean progeny count, averaging approximately 650 and 690 neonates, respectively. This output surpassed that of the parental genotypes (RR × RR and CC × CC), which produced average progenies of 400 and 520, respectively. Conversely, when reared on corn leaves diet, a divergent reproductive performance was observed among the hybrid populations. The CR × CR cross exhibited the most pronounced fecundity, with a mean progeny count of 550, while the RC × RC cross experienced a sharp decline in reproductive success, yielding a mean progeny count of approximately 120. The parental genotypes, in comparison, produced intermediate progeny counts, averaging 340 for RR × RR and 250 for CC × CC.

### Gene expression differences and downstream functional analysis

In our laboratory populations, survivability of rice mitotypes was too low on corn diet, so we performed both RNA-seq and DNA-seq on the same individuals from the rearing on Poitout diet only. We confirmed that parental strains genomes are different and almost homozygous, while reciprocal hybrids were heterozygous at all autosomal positions and clearly aligned on the PC1 axis of the principal component analysis (PCA) (**Supplementary Fig. S2-4, Fig. S8**). Examination of both whole genome sequencing and RNA-seq dataset by PCA showed a strong genotype effect on global transcriptional expression (**Supplementary Fig. S5**, **Fig. S8A**), with the hybrid genotypes being clearly separated from the parental strains on PC1. This effect is observed on autosomes (**Supplementary Fig. S8B**) as well as on sex chromosome (**Supplementary Fig. S8C-D**). Parental strains are separated along PC2 as well as reciprocal hybrids, even though they have similar genotypes. This could be indicative of a parent-of-origin effect. Additionally, our results show no definitive evidence of sex-related expression bias in the PCA dataset, particularly in relation to chromosome 29 (**Supplementary Fig. S8C-D**).

To identify the genes affected by the variations in genotypes, we first performed trajectory analysis of differentially expressed genes (DEGs) across the four FAW genotypes, with the objective of delineating genotype-associated gene expression profiles from the RNA-seq data. Then, we conducted classical pairwise comparisons between RC vs. CR to understand how parent-of-origin can modify the global expression of larval transcriptome of FAW.

#### Hybridization effect showed by trajectory clustering

Of the 10 clusters produced by our trajectory analysis (**Supplementary Fig. S15**), cluster 3 delineated 405 DEGs that were associated with both hybrids (**Fig. 2A**), which could be involved with the observed heterosis effect on the phenotypic results. The two hybrid genotypes (CR and RC) showed a congruent over-expression of these genes, whereas the parental strains (CC and RR) exhibited an under-expression.

**Figure 2.**
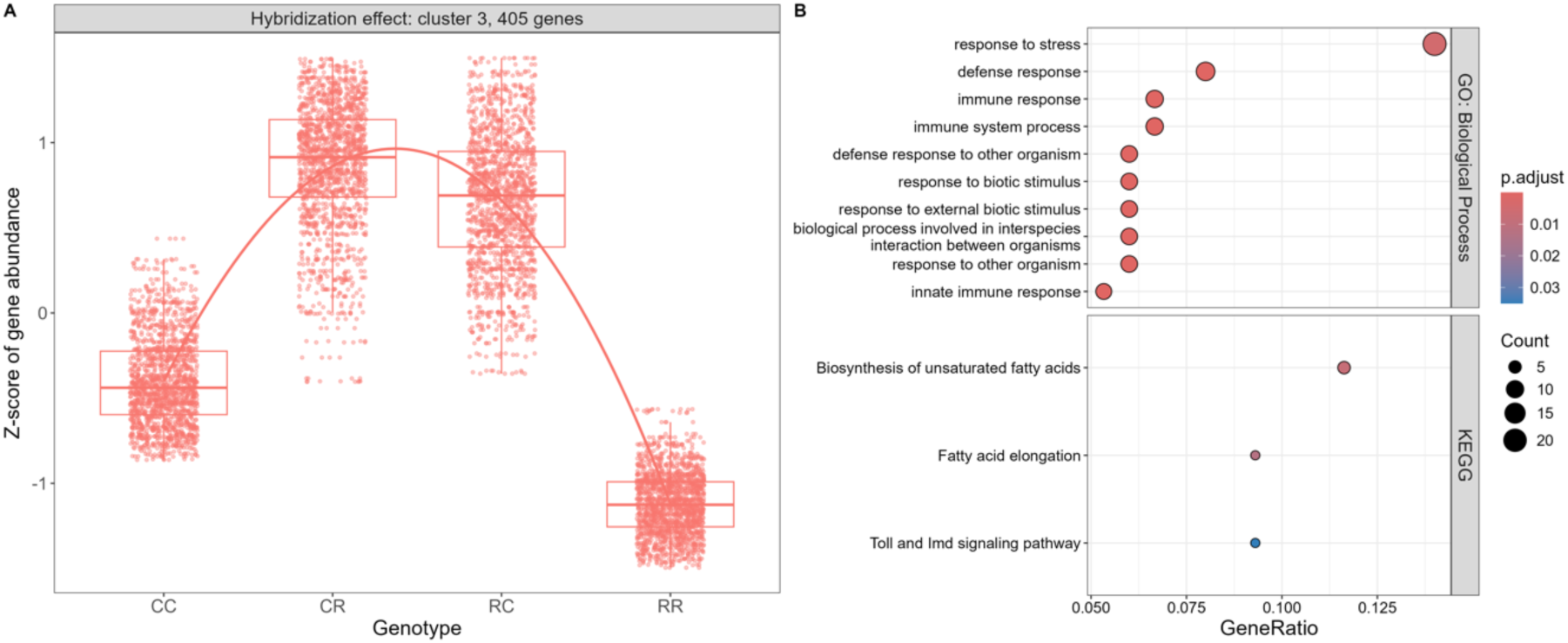
Selected cluster 3 of DEGs showing expression pattern of hybridization effect in the four FAW genotype transcriptomes and the related functional enrichment analysis. (A). On the left of the parallel coordinate plot, the cluster is labeled with 405 genes. (B). On the right side of the plot, the top 10 significantly enriched GO Biological Process terms and all the enriched KEGG pathways were highlighted. All gene expression clusters were given in Supplementary Figure **S15**.

Upon retrieval of all DEGs, an Over Representation Analysis (ORA) was conducted to identify enriched BP terms and KEGG pathways. The top 10 significantly enriched Biological Process (BP) terms in the hybrids compared to the parental strains were highlighted (**Fig. 2B**), with a particular emphasis on those associated with immune responses, such as “response to stress” (GO:0006950, adjusted *p*-value = 0.0045), “defense response” (GO:0006952, adjusted *p*-value = 1.75e-06), and “immune response” (GO:0006955, adjusted *p*-value = 1.35e-05), indicating a potential role of these processes in the hybrid vigor phenotypes. Furthermore, three enriched KEGG pathways were identified within the hybrid genotypes, which were “Biosynthesis of unsaturated fatty acids” (ko01040, adjusted *p*-value = 0.0080), “Fatty acid elongation” (ko00062, adjusted *p*-value = 0.0127), and “Toll and Imd signaling pathway” (ko04624, adjusted *p*-value = 0.035).

#### Candidate genes involved in hybridization effect in FAW

We then retrieved the genes associated with the top 10 enriched BP terms and the KEGG pathways. A total of 28 genes were identified in aggregate (as detailed in **Fig. 3** and **Supplementary Table S1**), showing indeed well-known immunity genes (Huot *et al*., 2020) such as *Spätzle*, *duox*, *cecropin*, *Spod-11-tox*, attacins and defensins.

**Figure 3.**
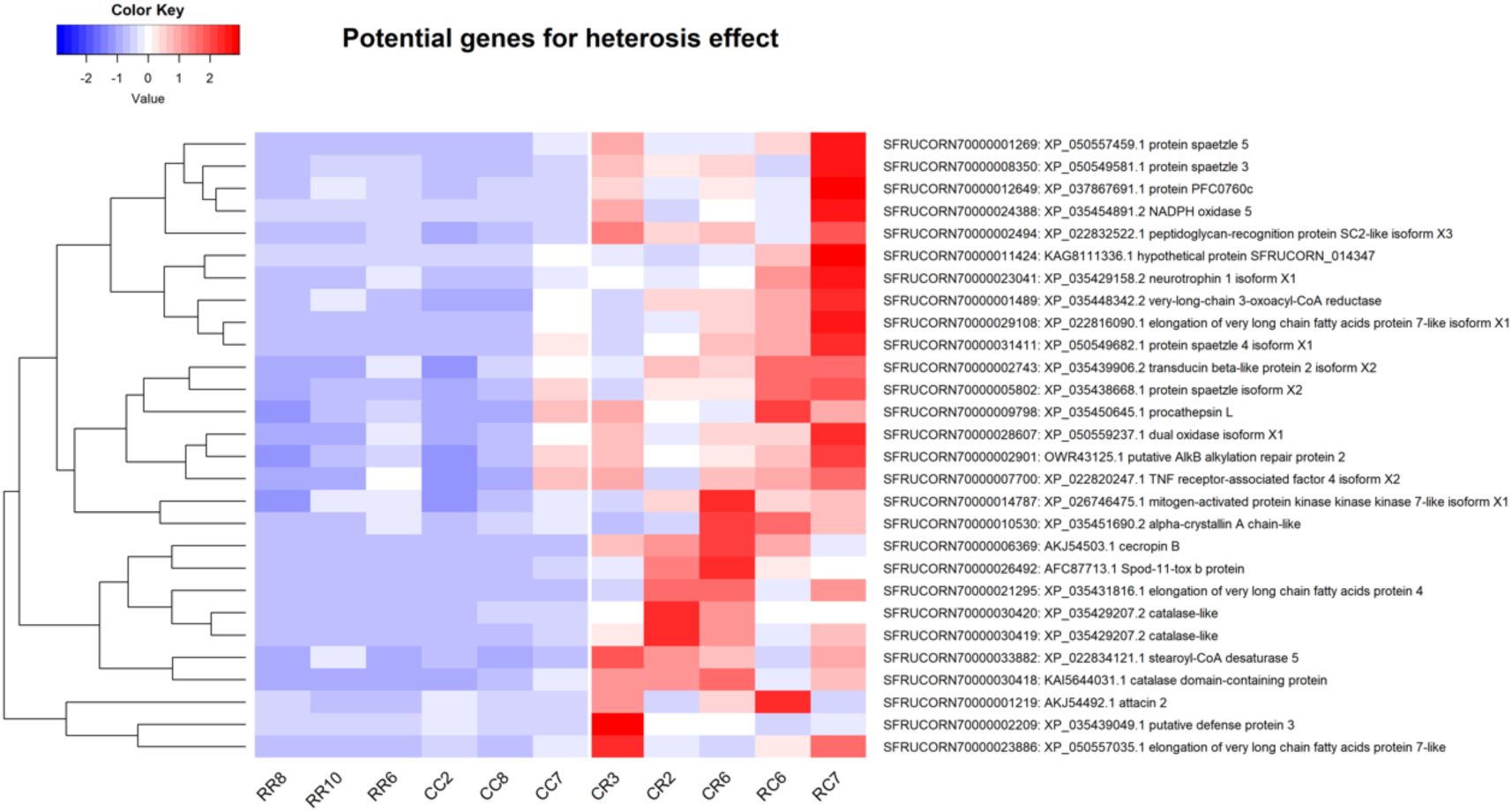
Heatmap showing the potential genes that might result in the heterosis effect of the hybrid genotypes (RC and CR) when compared to the parental strains (RR and CC) of FAW.

#### Maternal effect showed by trajectory clustering

As illustrated in **Figure 4A**, cluster 6 grouped 231 DEGs that were associated with the maternal effect in FAW. The genotypes with a rice strain maternal background, RR and RC, exhibited a consistent over-expression of these genes which were under-expressed in the genotypes with a corn strain maternal background, CC and CR. Following the retrieval of all these DEGs, an ORA was performed to identify enriched BP terms and KEGG pathways (**Fig. 4B**). While no KEGG pathways met the established threshold for significance, three enriched BP terms were identified, including “sulfur compound metabolic process” (GO:0006790, adjusted *p*-value = 0.0061), “cellular modified amino acid metabolic process” (GO:0006575, adjusted *p*-value = 0.030), and “glutathione metabolic process” (GO:0006749, adjusted *p*-value = 0.030).

**Figure 4.**
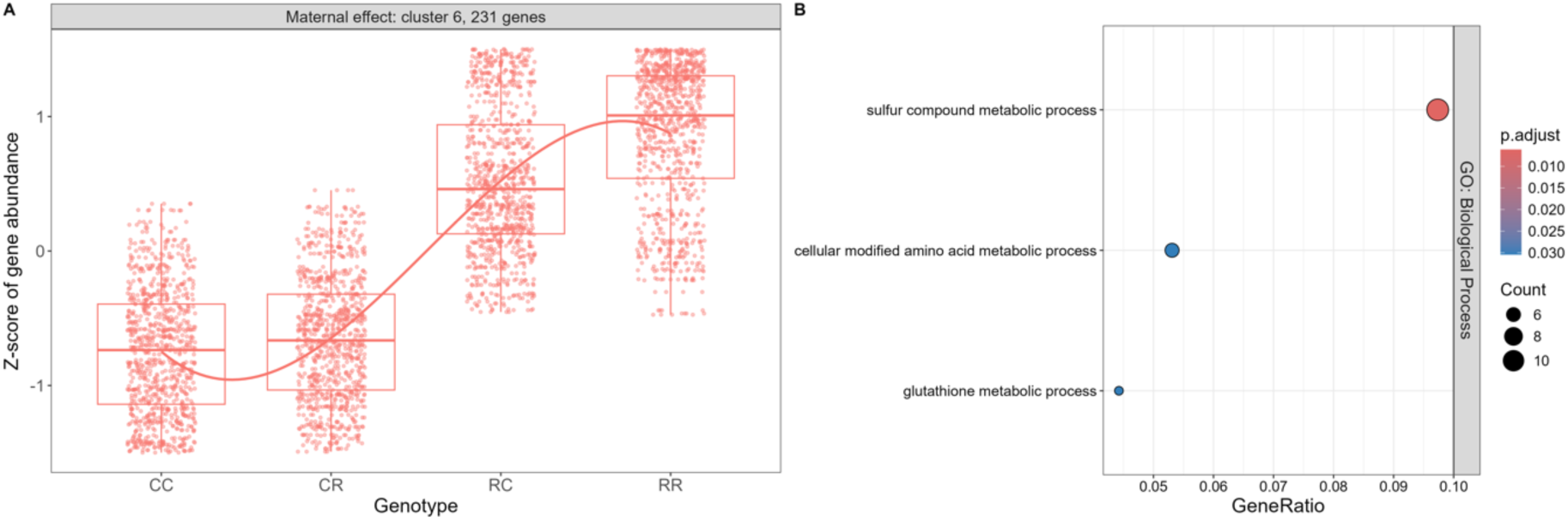
Selected cluster 6 of DEGs showing expression pattern of maternal effect in the four FAW genotype transcriptomes and the related functional enrichment analysis. (A). On the left of the parallel coordinate plot, the cluster is labeled with 231 genes. (B). On the right side of the plot, there were no enriched GO terms for the DEGs in this cluster, but three enriched KEGG pathways were highlighted. All gene expression clusters were given in Supplementary Figure **S15**.

#### Candidate genes involved in maternal effect in FAW

Subsequently, we extracted the genes associated with the three enriched BP terms to identify potential target genes that could be responsible for the maternal effect observed in FAW. A total of 12 genes were identified (as shown in **Figure 5** and **Supplementary Table S2**), including “5-oxoprolinase isoform X1”, “5-oxoprolinase-like”, “Bifunctional 3’-phosphoadenosine 5’-phosphosulfate synthase isoform X1”, “Putative fatty acyl-CoA reductase CG5065 isoform X2”, “Luciferin 4-monooxygenase”, “luciferin sulfotransferase-like”, “ATP-citrate synthase”, and “Trimethyllysine dioxygenase”, “Glutathione S-transferase-like”, “Sulfotransferase 1B1-like”, “Persulfide dioxygenase ETHE1”, and “Glutamate--cysteine ligase regulatory subunit”, which are associated with a few different metabolic processes.

**Figure 5.**
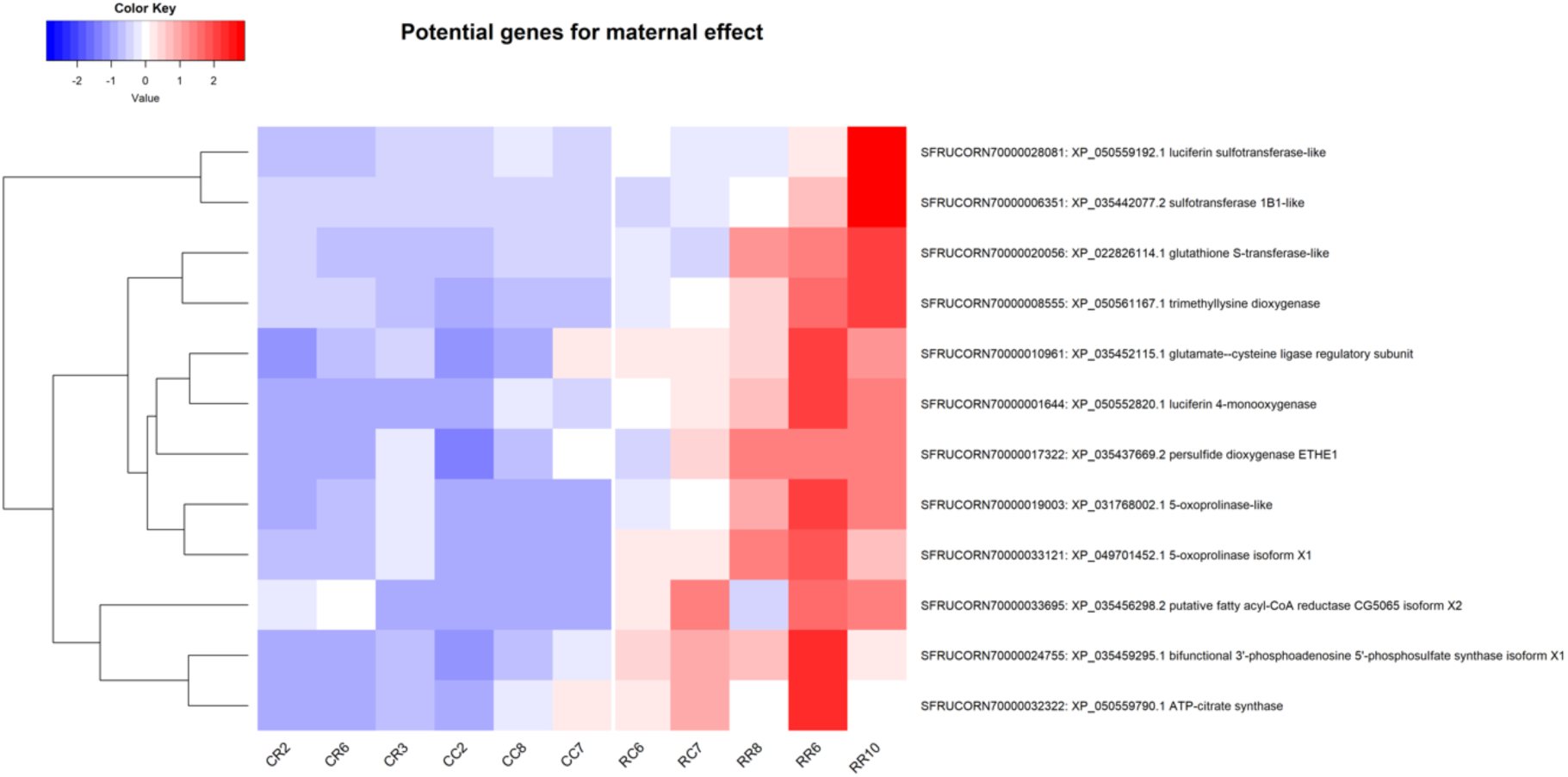
Heatmap showing the potential genes that might result in the maternal effect of the rice-strain maternal background genotypes (RC and RR) when compared to the corn-strain maternal background genotypes (CR and CC) of FAW.

#### Transcriptional differences between hybrid genotypes RC and CR

To evaluate the potential influence on gene expression patterns that may emerge from the interchange of genetic backgrounds between these genotypes, we compared the transcriptional profiles of the two hybrid genotypes, specifically RC versus CR, by a standard DESeq2 analysis (Wald test, see **Supplementary Method**).

While CR and RC hybrids have mostly the same genetic structure - they are heterozygous for all positions for males, except for the mitochondrial genome and the sex chromosomes for females - we observed a phenotypic difference between the CR and RC larvae (**Fig. 1**). This effect was particularly pronounced when reared on corn leaves (**Fig. 1D**), suggesting a parent-of-origin effect. PCA analysis of the RNA-seq data showed that the samples were grouped by hybrid (74% of explained variance on PC1; **Fig. 6A**). Differential gene expression analysis with DESeq2 identified 514 genes overexpressed in RC compared to CR, and 682 genes overexpressed in CR compared to RC (adjusted *p-*value < 0.05, **Fig. 6B**).

**Figure 6.**
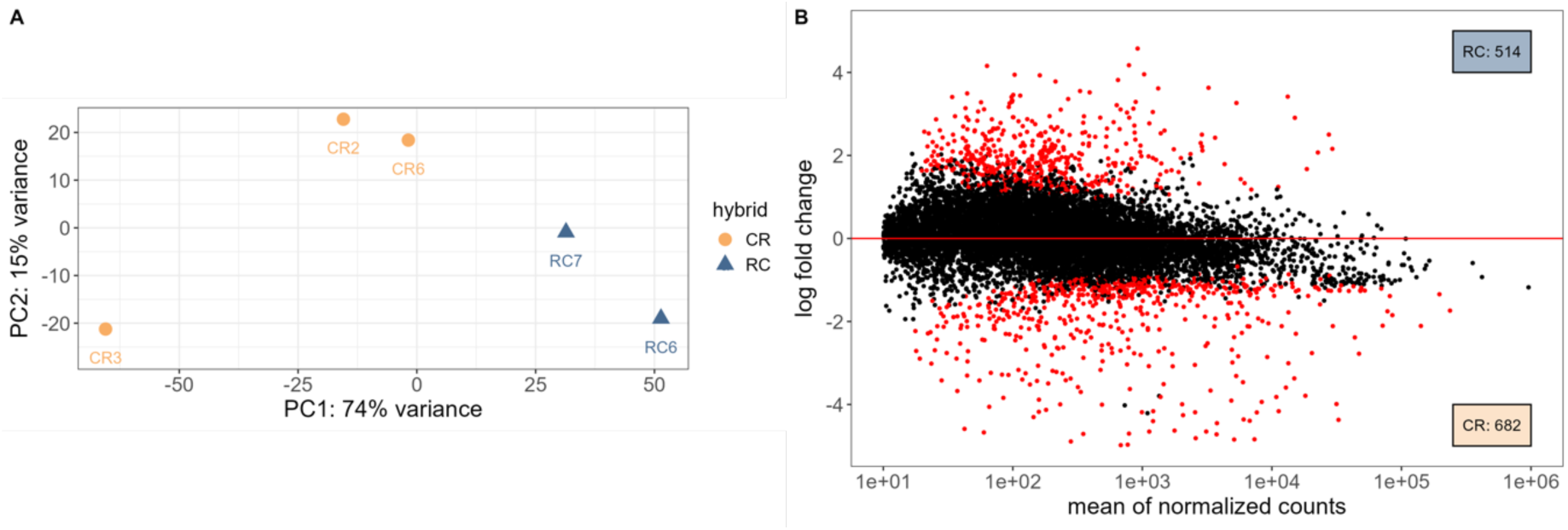
Transcriptional response of hybrid genotype (RC) versus the reciprocal hybrid genotype (CR). (A). Principal component analysis (PCA) on normalized RNA-seq reads for all the samples of RC and CR. (B). Multidimensional scaling plot (MA-plot) reporting the log2 fold changes between RC and CR over the mean of normalized counts. Each dot in the MA-plot signifies an individual gene, with non-significant differential expression represented by black dots, and genes exhibiting significant differential expression marked by red dots.

We retrieved all the DEGs of both RC and CR and performed ORA for GO and KEGG enrichment analysis (**Fig. 7, Supplementary Table S5-6**).

**Figure 7.**
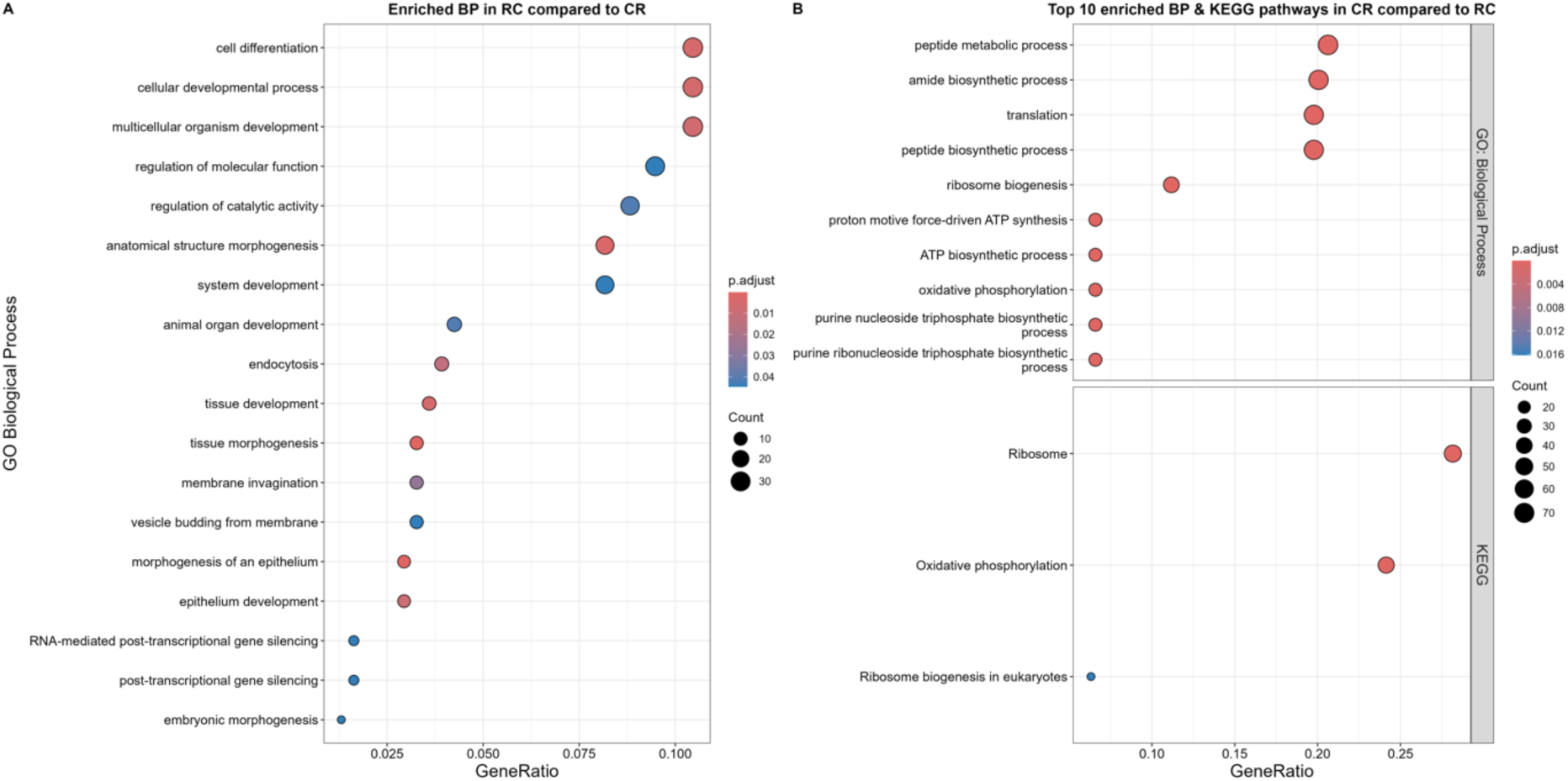
Enriched functional analysis of the DEGs for both RC hybrid and CR hybrid of FAW. (A). The enriched BP in RC compared to CC; (B). The top 10 enriched BP and all KEGG pathways in CR compared to RC.

For DEGs in RC hybrid, our analysis only yielded significantly enriched BP terms (**Fig. 7A**), the majority of which are associated with cellular processes and development, such as “cell differentiation” (GO:0030154, adjusted *p*-value = 0.0047), “cellular developmental process” (GO:0048869, adjusted *p*-value = 0.0047), and “anatomical structure morphogenesis” (GO:0009653, adjusted *p*-value = 0.0027). Intriguingly, we also revealed a subset of BP terms pertinent to gene silencing mechanisms, including RNA-mediated post-transcriptional gene silencing (GO:0035194, adjusted *p*-value = 0.045), and “post-transcriptional gene silencing” (GO:0016441, adjusted *p*-value = 0.045). Five genes were associated to “RNA-mediated post-transcriptional gene silencing” (GO:0035194) (**Supplementary Table S5**), including “putative helicase mov-10-B.1”, “endoribonuclease Dcr-1 isoform X1”, “endoribonuclease Dicer”, “protein Gawky isoform X1”, and “hypothetical protein SFRUCORN_019532 (HENMT1)” overexpressed in RC (**Fig. 8**). These genes are likely involved in the biogenesis and function of small RNAs, which mediate the silencing of target genes and transposable elements at the post-transcriptional level. Their overexpression may be triggered by inter-strain genomic incompatibilities within hybrids.

**Figure 8.**
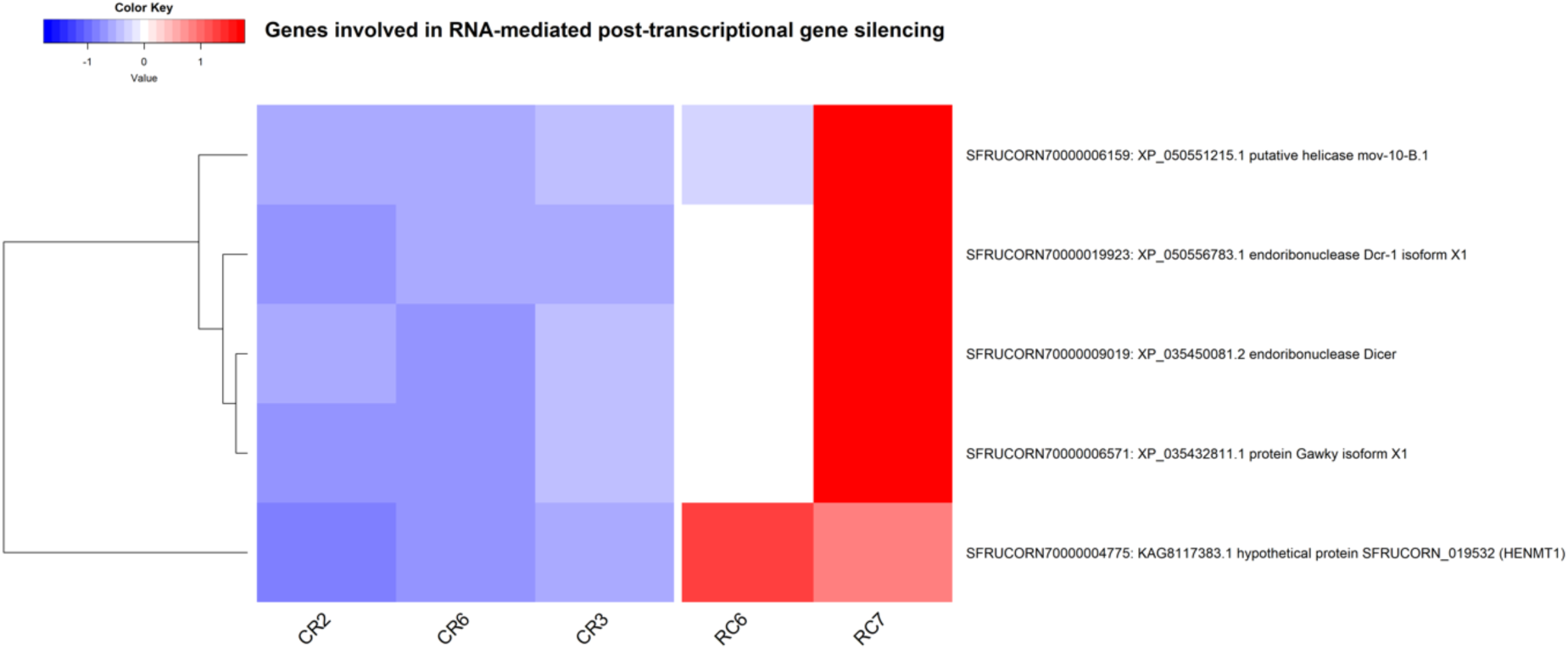
Heatmap showing the enriched genes of RNA-mediated post-transcriptional gene silencing in FAW RC hybrid compared to CR hybrid.

For CR hybrid DEGs, our enrichment analysis revealed BP terms associated with fundamental cellular activities such as translation, metabolic processes, biosynthesis, ribosome biogenesis, and energy production. The top 10 enriched BP terms (**Fig. 7B**) were “peptide metabolic process” (GO:0006518, adjusted *p*-value = 1.37e-23), “amide biosynthetic process” (GO:0043604, adjusted *p*-value = 2.74e-23), “translation” (GO:0006412, adjusted *p*-value = 7.86e-26), “ribosome biogenesis” (GO:0042254, adjusted *p*-value = 1.12e-15), “proton motive force-driven ATP synthesis” (GO:0015986, adjusted *p*-value = 7.85e-15), and “oxidative phosphorylation” (GO:0006119, adjusted *p*-value = 2.96e-14). For KEGG pathway enrichment, three significant pathways were identified, namely “Ribosome” (ko03010, adjusted *p*-value = 7.35e-31), “Oxidative phosphorylation” (ko00190, adjusted *p*-value = 7.69e-24), and “Ribosome biogenesis in eukaryotes” (ko03008, adjusted *p*-value = 0.016).

We identified 44 target genes from the “oxidative phosphorylation” process (**Fig. 9, Supplementary Fig. S16**), and most of these genes are crucial components of the mitochondrial electron transport chain and ATP synthesis, which are essential processes for cellular energy production. Specifically, there were several genes encoding subunits of key protein complexes, including NADH Dehydrogenase (Complex I), Cytochrome c Oxidase (Complex IV), Cytochrome b-c1 Complex (Complex III), ATP Synthase (Complex V), V-type Proton ATPase, and some other proteins.

**Figure 9.**
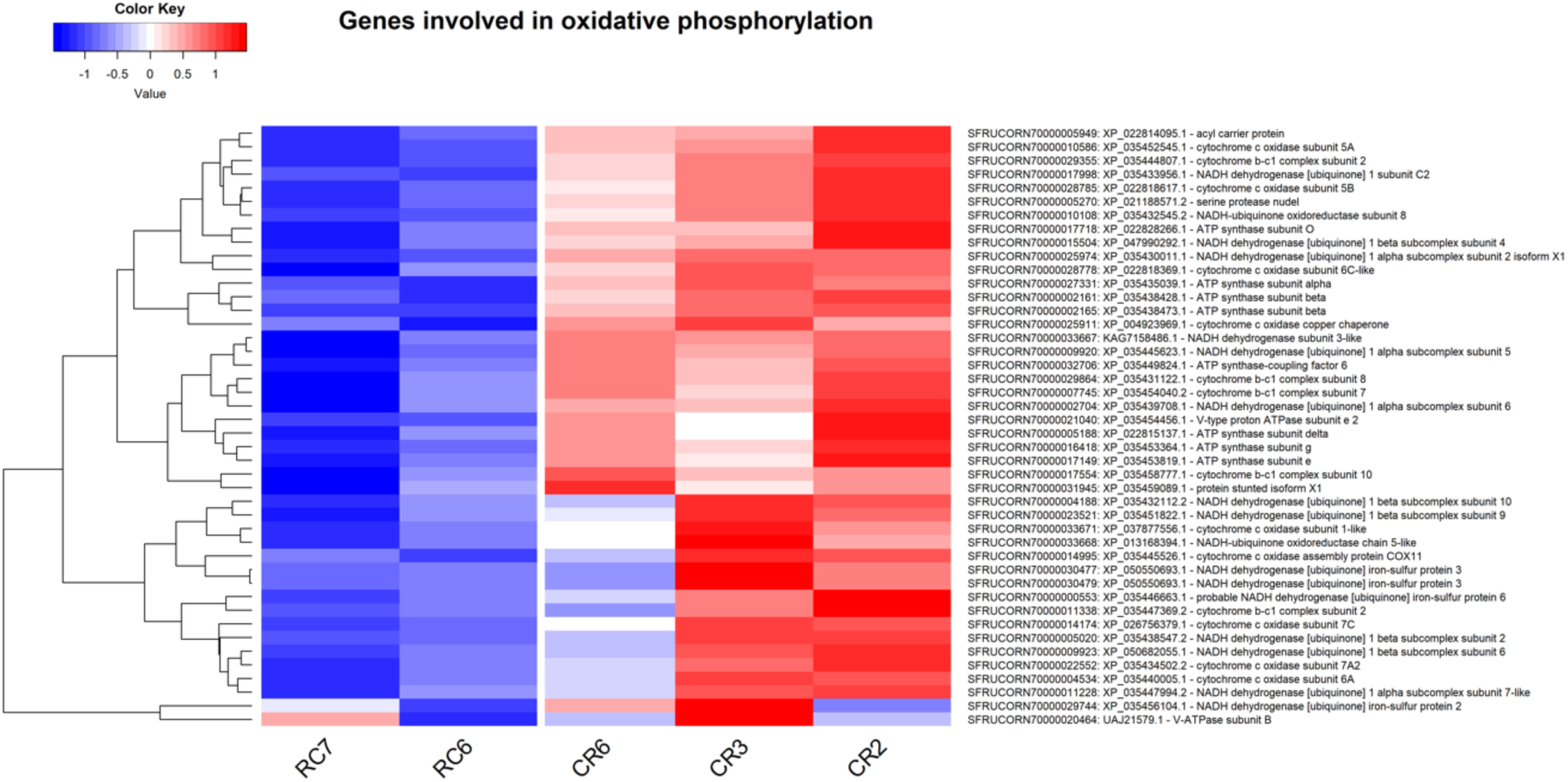
Heatmap showing the enriched genes of oxidative phosphorylation in FAW CR hybrid compared to RC hybrid.

## Discussion

The objective of this study was to identify the mechanisms explaining the pronounced and consistent transcriptional plasticity observed between the two FAW strains (Orsucci *et al*., 2022), which we hypothesized to underline the distinct plant adaptation capabilities of each strain. The most used genetic markers to differentiate FAW strains are localized on the mitochondrial DNA and sex chromosomes. Phylogeny of FAW based on mitochondrial DNA produces two clades, one of them clearly corresponding to the SfR host-strain (Kergoat *et al*., 2012) and Z-chromosome being the target of selective sweeps (Fiteni *et al*., 2022). To assess the potential impact of these host-associated loci on gene expression, we conducted a comparative analysis of larval performance in parallel with gene expression profiling of reciprocal hybrids between the FAW strains relative to the parental strains.

### Overall genotype effect on FAW larval performance

In our laboratory populations, we observed a strong incidence of cannibalism among FAW individuals when they were fed on corn plants. To mitigate this behavior in the current study, we conducted rearing with individual larvae instead of mass rearing (see **Methods**). Phenotypic measurements in these conditions were consistent with our prior findings (Lan and Nègre, 2023).

On both diets (especially on Poitout), we identified a heterosis effect on larval weight gain for both hybrids CR and RC, a phenomenon also reported in other research (Velásquez-Vélez *et al*., 2011). Heterosis, commonly known as hybrid vigor, is a multifaceted phenomenon where performance of the hybrid offspring is increased compared to their parental strains (Mather and Jinks, 1971; Virmani, 1994). Honeybee and the silkworm are examples of insects where the harnessing of heterosis has been particularly successful (Hoy, 1976).

Interestingly, when reared on a corn plant diet, a pronounced maternal effect was observed, with genotypes possessing a rice-strain maternal lineage (RR and RC) exhibiting significantly lower survival rates compared to those with a corn-strain maternal background (CC and CR) (**Fig. 1-D**). In assessing the reproductive capacity of the hybrid populations (RC × RC and CR × CR), we observed a significant increase in mean progeny count, particularly when fed on Poitout artificial diet (**Supplementary Fig. S1-A**). In contrast, on a corn plant diet, the RC × RC population displayed diminished fecundity (**Supplementary Fig. S1-B**). Nutritional intake is crucial in the life histories of insect species, with diet quality and quantity exerting a profound influence on their reproductive success (Fischer *et al*., 2004; Bauerfeind and Fischer, 2005; Kehl and Fischer, 2012). The reduced fecundity of the RC × RC population on corn plants could be attributed to suboptimal nutritional conditions or physiological incompatibilities with this host plant. Notably, the CR × CR hybrid population produced the greatest number of offspring on both diets, suggesting a robust adaptive potential for this genotype. These results indicate that the CR × CR hybrid may possess a genetic advantage that enhances its reproductive success in varied environmental conditions. We previously hypothesized that dominance at heterozygous loci in loci may explain enhanced metabolism during larval stage (Lan and Nègre, 2023). We thus performed RNA-seq on larvae from different genotypes in the same rearing conditions to verify whether heterozygosity can affect the expression of particular sets of genes.

### Overall genotype effect on FAW transcriptional gene expression

The PCA of RNA-seq data showed that the observed clustering was predominantly determined by genotype (**Supplementary Fig. S8**). PC1 differentiated the hybrid genotypes, CR and RC, from the parental genotypes, CC (except for CC7) and RR, suggesting a hybridization effect, while PC2 segregated individuals along maternal lineages, hinting at a possible maternal effect on gene expression profiles (**Supplementary Fig. S8C-D**). Through our trajectory analysis of the DEGs identified by the likelihood ratio test (LRT) (see **Supplementary Method**), we grouped subsets of genes associated with each genotype (**Supplementary Fig. S15**).

#### Heterosis effect in FAW

In our previous studies, the heterosis effect in FAW hybrids was meticulously documented across a range of phenotypic traits through mass rearing method (Lan and Nègre, 2023). Our current investigation, conducted on individual rearing, has revealed a consistent enhancement in larval weight among FAW hybrids (**Fig. 1**). This improvement was observed in both artificial diet and natural corn diet (**Fig. 1A-B**). Through a comprehensive clustering and functional enrichment analysis, we identified a list of DEGs associated with an over-expression in hybrid genotypes (**Fig. 2** and **Supplementary Table S1**) dominated by immune-related genes, including “Spod-11-tox b protein”, “Attacin 2”, and “Cecropin B”, which are integral effector components of the immune response and defense mechanisms in FAW (Legeai *et al*., 2014).

In addition, we detected genes involved in lipid metabolism, such as “Elongation of very long chain fatty acids protein 7-like isoform X1”, which suggests a genetic basis for the modulation of energy storage and utilization in the hybrid genotypes. This metabolic flexibility may be particularly advantageous for the hybrids, allowing them to efficiently allocate resources in response to varying environmental conditions and nutritional availabilities (Kehl and Fischer, 2012; Park *et al*., 2016; Zhang *et al*., 2019). However, it is well-known that a trade-off exists in insects between the production of energy and the production of immunity factors by the fat body (Gupta *et al*., 2022). Hybrids appear to overcome or lessen the typical trade-off observed in non-hybrid genotypes, achieving a more refined balance between metabolism and immunity. This optimized allocation of resources may contribute to hybrid vigor (heterosis) in FAW.

#### Maternal effect in FAW

Transcriptional response of herbivores to their host plants can be shaped by the host plant experiences of previous parental generations (Lillycrop *et al*., 2008; Müller *et al*., 2017). When we conducted a trajectory analysis on our RNA-seq dataset, we noticed a strong maternal effect on gene expression of FAW, with more than 200 genes overexpressed in the rice-mother maternal background genotypes (RC and RR) compared to the corn-mother maternal genotypes (CC and CR). Through the functional enrichment analysis, we identified a significant overrepresentation of genes related to a few metabolic processes. Indeed, research has indicated that FAW engages in active metabolic processes throughout its lifespan, which are crucial for its physiological functioning and development (Wang *et al*., 2021). Notably, during its larval stage, FAW is known to undergo detoxification and xenobiotic metabolism, processes that are essential for managing exposure to foreign substances and potentially harmful compounds.

In our investigation of the maternal effects in FAW, we identified several candidate genes that may contribute to these phenotypic influences. One such gene of interest is the enzyme luciferin 4-monooxygenase. Luciferase, while renowned for its luminescent function, also catalyzes the synthesis of fatty acyl-CoA via a biochemical mechanism (Oba *et al*., 2003; Mofford *et al*., 2014). We have detected an additional target gene, fatty acyl-CoA reductase, which is implicated in the biosynthesis of precursors for pheromones (Wicker-Thomas *et al*., 2015) and cuticular hydrocarbons (CHCs) (Blomquist and Bagnères, 2010; Holze *et al*., 2021). CHCs primarily offer a critical barrier against desiccation, and interestingly, they have also been demonstrated to possess pheromonal properties, playing a crucial role in enhancing mating success (Jallon and Wicker-Thomas, 2003; Howard and Blomquist, 2005). Oenocytes, recognized as specialized organelles for the biosynthesis of CHCs, rely on lipid metabolism within adipose tissue to produce pheromones (Chung *et al*., 2014; Wicker-Thomas *et al*., 2015). The maternal transfer of these biochemical precursors may affect chemical signaling and the development of the cuticular structure in progeny, potentially impacting their adaptation and survival. Indeed, the pheromone composition of FAW hybrid female has been found to exhibit a pattern of maternal inheritance for some major pheromone components, suggesting a genetic basis for these traits (Groot *et al*., 2008). Besides, we noted similarities in the FAW external morphological characteristics of the larval cuticle, particularly between RR and RC, and likewise between CC and CR individuals (Lan and Nègre, 2023).

### Hybrid effect on FAW gene expression

Through pairwise comparison, DEGs overexpressed in CR compared to RC genotypes, are massively associated with “translation” and with “oxidative phosphorylation” (**Supplementary Table S6**). This metabolic pathway was not only enriched in our GO analysis but also prominently featured in the KEGG enrichment analysis when examining CR versus RC (**Fig. 8B, Supplementary Fig. S16**), reflecting the higher mitochondrial output of the C-strain that we hypothesize. We have pinpointed 44 key target genes depicted in **Figure 9**. These genes, being central to the mitochondrial electron transport chain and ATP synthesis, could be the molecular basis for the CR genotype’s superior performance compared to RC. Their involvement in oxidative phosphorylation suggests that they may confer an enhanced capacity to withstand oxidative stress, a common challenge for cells undergoing high metabolic activity. The ability to maintain efficient energy production under stress could be a key factor in the CR’s resilience. Moreover, the upregulation of these genes might also be linked to an improved antioxidant defense system, which is essential for neutralizing reactive oxygen species (ROS) that can damage cellular components (Hong *et al*., 2024). By having a more robust defense mechanism, the CR hybrid may be better equipped to mitigate the harmful effects of ROS, thereby maintaining cellular homeostasis and functionality. Furthermore, the genes’ roles in energy metabolism could indirectly influence other defense mechanisms. For instance, sufficient ATP supply is often required for the activation of immune responses and the upregulation of stress-related proteins (Pearce and Pearce, 2013; Buck *et al*., 2015). Thus, the CR hybrid’s heightened performance might be attributed to a combination of efficient energy production and a reinforced defense system and contribute to the observed heterosis effect in FAW.

Among the enriched BP terms in RC compared to CR, we noticed two that are related to gene silencing mechanisms, namely “RNA-mediated post-transcriptional gene silencing” (GO:0035194), and “post-transcriptional gene silencing” (GO:0016441). We further extracted the DEGs behind the first BP term and detected a few target genes, including MOV-10 (Moloney leukemia virus 10), two Dicer, Gawky, and HENMT1 (HEN1 methyltransferase homolog 1). These genes collectively contribute to the RNA-mediated gene silencing mechanisms, particularly in the RNA interference (RNAi) pathway and the PIWI-interacting RNA (piRNA) pathway. Specifically, RNAi could involve the processing of mRNAs by endoribonucleases such as Dicer, resulting in the production of small interfering RNAs (siRNAs) (Bernstein *et al*., 2001). The putative helicase MOV-10 might then assist in the unwinding and incorporation of these siRNAs into the RNA-induced silencing complex (RISC) (Meister *et al*., 2005). Within RISC, the protein Gawky could play a role in mRNA decapping and degradation, further enhancing gene silencing (Rehwinkel *et al*., 2005; Schneider *et al*., 2006). Additionally, the hypothetical protein HENMT1 might contribute to the stabilization of piRNAs through methylation, ensuring the effective regulation of transposable elements (TEs) and other RNA targets to maintain genome integrity (Gainetdinov *et al*., 2021; Haase, 2022). Indeed, comparative analysis between the RR and CC strains has revealed a higher prevalence of TEs in the CC strain, a finding supported by Orsucci *et al*. (2022). Our dataset corroborates this observation, showing an enrichment of TEs in the CC strain when compared to RR strain (**Supplementary Table S4**), specifically reverse transcriptases and the putative harbinger transposase-derived nuclease 1 (HARBI1). TEs are known to induce F_1_ hybrid incompatibility and intrastrain dysgenesis in certain *Drosophila* species, as reported by Castillo and Moyle (2022). The pivotal role of the piRNA pathway in the antiviral immune response has been investigated in the Lepidopteran cell lines of both *Bombyx mori* and *Trichoplusia ni* (Santos *et al*., 2022), as well as in mosquitoes (Hess *et al*., 2011; Léger *et al*., 2013). Virus infection may also trigger a piRNA-based immune defense in the fat body of FAW (Robin *et al*., 2023). It is plausible that the genetic cross between the two FAW strains has activated the gene silencing pathway in RC due to the deregulation of TEs in the different genotypes mimicking the presence of viral elements.

## Conclusion

The goal of our study was to identify the genetic factors responsible for the phenotypic variation and consistent transcriptional divergence observed between FAW strains. We have pinpointed several potential genetic determinants that may account for these transcriptional differences and have profiled their expression during the larval stage. Additionally, we have identified a plethora of candidate genes associated with the heterosis and maternal effects influencing larval performance and overall gene expression, which could be key to understanding the invasive nature and host-plant adaptation in FAW. From our broad overview of transcriptional patterns, subsequent studies should investigate the functional roles and regulatory mechanisms of these candidate genes at physiological and cellular levels.

## Acknowledgments

This work was supported by funding from SPE department of INRAE for NN and by the scholarship from China Scholarship Council (CSC) under the collaborative program of CSC and Agreenium (CSC NO. 202008440426; https://en.agreenium.fr/page/agreenium) for LL. We thank the quarantine insect platform (PIQ) for providing the infrastructure needed for insect pest experiments. We are grateful to Gaëtan Clabots, Raphaël Bousquet, and Dylan Valenza for maintaining the insect collections of the DGIMI laboratory in Montpellier, France. We appreciate the help from Allyson Moureaux for producing the BAI files and sorted BAM files of the RNA-seq dataset.

