## Supplementary Material for "Parent-of-origin effect on global gene expression and host-plant adaptation in *Spodoptera frugiperda* (Lepidoptera: Noctuidae)"

running title: Gene expression differences in fall armyworm hybrids

**Laijiao Lan^1^**
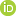
**^<https://orcid.org/0000-0003-0406-2199>^, Stéphanie Robin^2,3^**
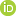
**^<https://orcid.org/0000-0001-7379-9173>^, Kiwoong Nam^1^**
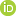
**^<https://orcid.org/0000-0003-3194-8673>^ and Nicolas Nègre^1^***
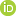
**^<https://orcid.org/0000-0001-9727-3416>^**

^1^ DGIMI, Univ Montpellier, INRAE, Montpellier, France

^2^ BIPAA, IGEPP, INRAE, Institut Agro, University of Rennes, Rennes, France

^3^ IRISA, University of Rennes, INRIA, CNRS, IRISA, Rennes, France

### Extended Scripts for Materials and Methods

**RNA-seq analysis pipeline**

*Differential gene expression analysis* **–** Differentially expressed genes (DEGs) were identified using the R package DESeq2 (v1.42.0, Love *et al.*, 2014), with pre-filtering to keep only rows that have a count of at least 10 for a minimal number of samples and an adjusted *p-value* < 0.05. The adjustment method for *p*-value is Benjamini-Hochberg (BH), which controls the false discovery rate, the expected proportion of false discoveries amongst the rejected hypotheses (Benjamini and Hochberg, 1995). Fold change shrinkage was applied on the count values with “apeglm” method (Zhu *et al.*, 2019) for downstream visualization.

*Pairwise Comparisons* – To discern variations in gene expression across genotype groups—especially RC vs. CR (and RR vs. CC as supplementary results)—DESeq2’s Wald test was employed. This statistical method involves doubling the tail integrals from the normal distribution to execute a two-tailed test. Consequently, genes with an adjusted *p-value* less than 0.05 post this adjustment were deemed significant within each pairwise comparison, aligning with a significance threshold (α) of 0.05.

*Model Comparisons* – To identify gene expression patterns that are significantly associated with different genotypes, the likelihood ratio test (LRT) statistic was applied through the DESeq2 package. We used a full model containing genotype of each sample and generated an intercept-only reduced model (reduced = ~ 1) to test the genotype effects on gene expression. Applying this methodology, genes exhibiting adjusted *p-value* below 0.05 were recognized as statistically significant, which corresponds to a significance level (α) of 0.05.

*Clustering* – The R package DEGreport (v1.40.0, Pantano, 2024) was utilized to conduct hierarchical clustering on vst-transformed counts of DEGs (adjusted *p* < 0.05) identified by the LRT for calculating pairwise gene expression among genotype conditions using Kendall’s rank. Hierarchical clustering is a technique that establishes a sequence of hierarchical relationships among data points, progressively dividing them into clusters to form a dendrogram. This visual representation illustrates the degree of similarity between data points. The “degPatterns” function within DEGreport is employed to segment the dendrogram at a height equivalent to the divisive coefficient of the clustering. This coefficient serves as a metric for the clustering structure within the dataset, effectively reducing the number of clusters for datasets with a more cohesive clustering pattern. By applying this approach, clusters of genes that exhibit analogous expression trajectories were created, and Z-scores of these genes were visualized. These gene clusters were then further analyzed through enrichment analysis to uncover biologically meaningful patterns and pathways associated with the observed expression changes.

*Enrichment analysis (GO and KEGG) –* Functional enrichment analysis plays a crucial role in deciphering the biological themes that are most representative of the complex datasets generated by high-throughput sequencing. To gain a deeper understanding of the roles of DEGs across the four genotypes for both the pairwise comparisons and the model comparisons, we initiated our investigation with Gene Ontology (GO) enrichment analysis. GO is a systematic framework that categorizes gene functions and delineates the interrelationships among these functions. It encompasses three primary domains: Molecular Function (MF), which pertains to the activities of the gene products; Cellular Component (CC), which relates to the parts of the cell where the gene products are active; and Biological Process (BP), which involves the pathways and functions of gene products in the context of an organism’s life. In our study, we concentrated primarily on the BP category to uncover the biological processes that were significantly enriched in each of the genotype comparisons. We applied blast2GO (v1.5.1) Command Line (Conesa *et al.*, 2005) for sequence alignment via sequence alignments (BLAST), identifying potential homologs for each input sequence and mapping them to GO terms. We set stringent criteria for the GO annotation process, applying an E-Value-Hit-Filter threshold of 1.0E-6 and a similarity cut-off of 55%, ensuring an evaluated functional annotation of the query sequences (Götz *et al.*, 2008). This meticulous approach allowed us to identify the most characteristic BP of the DEGs associated with each genotype.

To check the biological pathways associated with the enriched gene sets, we expanded our analytical scope to include the Kyoto Encyclopedia of Genes and Genomes (KEGG) pathway analysis. KEGG is a widely recognized database that provides a comprehensive collection of graphical representations of biological pathways, which are crucial for understanding the molecular interactions and functions within cells. By integrating KEGG pathway analysis into our study, we aimed to identify the specific biological pathways that are significantly impacted by the gene expression changes observed across different genotypes. This approach allows us to map the DEGs onto known pathways, contributing to the discovery of coordinated gene functions and the elucidation of the molecular mechanisms that drive the phenotypic differences associated with the genotypes. To obtain a background data set of the KEGG annotation information, 21 species (*Drosophila melanogaster* (fruit fly), *Musca domestica* (house fly), *Anopheles gambiae* (malaria mosquito), *Aedes aegypti* (yellow fever mosquito), *Apis mellifera* (honey bee), *Bombus terrestris* (buff-tailed bumblebee), *Solenopsis invicta* (red fire ant), *Camponotus floridanus* (Florida carpenter ant), *Polistes canadensis*, *Nasonia vitripennis* (jewel wasp), *Microplitis demolitor*, *Tribolium castaneum* (red flour beetle), *Bombyx mori* (domestic silkworm), *Danaus plexippus* (monarch butterfly), *Plutella xylostella* (diamondback moth), *Trichoplusia ni* (cabbage looper), *Helicoverpa armigera* (cotton bollworm), *Acyrthosiphon pisum* (pea aphid), *Aphis gossypii* (cotton aphid), *Daphnia pulex* (common water flea), and *Tetranychus urticae* (two-spotted spider mite)) were considered as background sets to annotate the best alignment sequences against our annotated genome (OGS7.1_20230705_proteins.fa) by using the KEGG database, which was accomplished by the KEGG Automatic Annotation Server (KAAS, <http://www.genome.jp/tools/kaas>) (v2.1, Moriya *et al.*, 2007). The KEGG Ortholog assignments and pathway maps were obtained using the bidirectional best hit method (BBH) on KAAS.

Further enrichment analysis was performed on DEGs based on the GO terms and KEGG pathways using clusterProfiler package (v4.10.0, Wu *et al.*, 2021) in R. This comprehensive tool enabled a detailed examination of the DEGs in the context of GO terms and KEGG pathways. We distinguished DEGs that were upregulated (log2 fold change > 0) from those that were downregulated (log2 fold change < 0) across each pairwise genotype comparison. These gene sets were independently subjected to Over Representation Analysis (ORA) (Boyle *et al.*, 2004). ORA assists in identifying whether the DEGs are significantly overrepresented within a particular biological category or pathway. We set stringent criteria for identifying enriched GO and KEGG pathways in ORA, using a *p*-value cutoff of 0.05 and a q-value cutoff of 0.2 with the universal “enricher” function within clusterProfiler. GO with > 0.7 similarity are removed to reduce redundancy among enriched GO terms using the “simplify” function in clusterProfiler. This approach ensured that only the most statistically significant and biologically relevant pathways were considered. Furthermore, we applied clusterProfiler to perform ORA analyses on the gene clusters that were previously generated using the DEGreport package (v1.40.0, Pantano, 2024). This dual application of clusterProfiler and DEGreport allowed us to comprehensively dissect the functional roles and pathways associated with the observed gene expression patterns, providing a robust framework for understanding the biological underpinnings of the genotype-specific gene expression profiles.

*Visualization* – To do quality check for the samples, PCA plots were generated with vst-transformed counts by using the plotPCA function in DESeq2 (v1.42.0, Love *et al.*, 2014). To visualize the DEGs identified using the methods described above, MA plots were produced with plotMA function in DESeq2. Both PCA and MA plots were further customized ggplot2 (v3.5.1, Wickham, 2016) for better visualization. Enriched genes after enrichment analysis were retrieved for each comparison (including the gene clusters generated from DEGreport) and were used to produce heatmaps using scaled raw counts to visualize the potential target genes, with the R package gplots using scaled raw counts (v3.1.3.1).

The overall RNA-seq analysis workflow can be seen in **Supplementary Figure S14**. Specifically, the complete annotated transcriptome was subjected to a differential expression analysis employing LRT in the DESeq2 package (v1.42.0, Love *et al.*, 2014). This LRT was utilized to evaluate the correlation between genotype and gene expression levels across all experimental groups based on their vst-transformed read counts, resulting in the identification of 2,024 DEGs with an adjusted *p*-value < 0.05. Subsequently, gene expression trajectories were analyzed across the distinct FAW genotypes using these 2,024 DEGs. This analysis was accomplished through divisive hierarchical clustering with the R package DEGreport (v1.40.0, Pantano, 2024). **Supplementary Figure S15** illustrates the comprehensive expression patterns of the 10 resultant clusters. We have highlighted two clusters that exemplify the characteristic expression patterns observed between hybrid genotypes and parental strains (**Fig. 2**), as well as between genotypes associated with corn and rice maternal backgrounds (**Fig. 4**). Additionally, we also presented two clusters that exhibited the differential expression patterns between the two strains (Supplementary **Fig. S9**). Furthermore, the Wald test was employed to assess the statistical significance of transcript count variances for each gene between each pairwise genotype comparison, namely RC vs. CR (also RR vs. CC as extended results, Supplementary **Fig. S10**-**S13**).

Both the DEGs within the selected clusters and the results of the pairwise comparisons underwent ORA using the clusterProfiler R package (v4.10.0, Wu *et al.*, 2021). This analysis aimed to identify enriched BP and KEGG pathways (ranked by the adjusted *p*-value with BH, FDR < 0.05), focusing on genes that exhibit dynamic regulation in a pattern consistent across the various FAW genotypes. Ultimately, genes associated with the enriched BP and KEGG pathways identified in each comparative analysis were extracted. Scaled raw counts of these genes were then utilized as the input to construct a heatmap with the R package gplot2, facilitating the visualization of differential gene expression patterns between the groups.

**Data availability**

.fastq files and .bam files for the RNA-seq dataset can be accessed publicly through the ArrayExpress collection at the following link:

https://www.ebi.ac.uk/biostudies/arrayexpress/studies/E-MTAB-14826

**Processed files for RNA-seq anaysis and R scripts repository**

Analysis files for this publication can be found on Zenodo at the following link: 10.5281/zenodo.14856499

### Extended Results

**Genotype confirmation**

To make sure that the two strain genomes analyzed are indeed different and each individual sample was of the correct genotypes before performing transcriptome analysis on individual larvae, both DNA and RNA of the same individuals from each genotype were extracted and purified at the same time (see **DNA and RNA extraction**). We first examined DNA sequencing alignments onto the mitochondrial genome, - represented in our reference genome by HiC_scaffold_320. Since our reference genome has been obtained from corn strain individuals, reads originating from corn strain should exhibit minimal to no single nucleotide polymorphisms (SNPs) when aligned to the reference genome. We thus could easily verify that the two laboratory strains we have used exhibit different mitochondrial genomes, with all individuals from corn-strain motherline (CC and CR) exclusively inherit a mitogenome associated with the corn strain, and that all individuals from rice-strain motherline (RR and RC) show a mitogenome characteristic of the rice strain. This can be visualized by observing the alignment .bam files in Integrative Genomics Viewer (IGV, Robinson *et al.,* 2011), (Supplementary **Fig. S2**), showing no SNP for the corn-strain mother individuals.

To further confirm the specific genotypes, we looked at the *Tpi* gene, which is often used for genotyping FAW strains (Nagoshi, 2010). This gene is located on the HiC_scaffold_29, which in our genome assembly, the *Spodoptera frugiperda* corn assembly v7.0 (Fiteni *et al.*, 2022), represents the Z chromosome. Again, all CC and RR individuals show a single genotype on *Tpi* (Supplementary **Fig. S3**), corresponding to their respective strain (Nagoshi, 2010). This again confirms that our laboratory strains contain only one representative haplotype for the Z chromosome. Hybrids should be heterozygous if they are male or have the Z haplotype of their father if they are female. Thus, CR2 should be a male Z_C_Z_R_, while CR3 and CR6 should be female since they have only one Z_R_ haplotype. Similarly, RC6 is a Z_R_Z_C_ male and RC7 a female with a Z_C_ chromosome. However, RC4 has only one Z_R_ haplotype, which should not be possible if it is a hybrid. Indeed, on the autosomes, for example on HiC scaffold_1 (Supplementary **Fig. S4**), CC and RR individuals contain only one genotype, while all hybrids, except RC4 are heterozygous, strongly suggesting that RC4 is a pure RR individual. This sample has thus been removed from subsequent RNA-seq analysis.

**Genetic homozygosity analysis of FAW genotypes**

With DNA-seq dataset, we also conducted the principal component analysis (PCA), which shows whole-genome differentiation between corn strain CC and rice strain RR (**Supplementary Fig. S5**). CR and RC hybrids were observed between the two parental strains, as expected if the genotypes of the hybrids are intermediate between their parents.

We also compared heterozygosity among the four FAW genotypes based on the number of heterozygously called positions (**Supplementary Fig. S6**). The average number of heterozygous positions in CC and RR ranged from 799 Kb to 957 Kb and 646 Kb to 730 Kb, respectively, which were lower than in CR (3,052 Kb to 3,332 Kb) and RC (3,127 Kb to 3,230 Kb) hybrids. Considering that the assembly size is 385,049 Kb, the heterozygosity of CC and RR is 0.20% – 0.24% and 0.16% – 0.18%, respectively. This heterozygosity corresponds to only 8.51% – 10.2% of sfC and 6.70% – 7.53% of sfR estimated from natural populations in Mississippi, where the average heterozygosity is 2.35% and 2.39%, respectively (Nam *et al.*, 2020).

**Effects of sex on widespread gene expression**

Since caterpillars have not developed mature sexual features, we should not expect a major influence of sex on gene expression. Nevertheless, we utilized RNA-seq coverage to assess the expression levels of the gene *Masculinizer* (*Masc*), which is located on the Z chromosome. This gene was meticulously re-annotated by BLAST with the GenBank sequence LC716474.1, representing the *Masc* complementary DNA (cDNA) in Sf9 cell line, which is derived from tissues of FAW, against the *Spodoptera frugiperda* corn assembly v7.0. The *Masc* gene is subject to repression by the PIWI-interacting RNA (piRNA) pathway in females, resulting in its exclusive expression in males (Katsuma *et al.*, 2015; Sakai *et al.*, 2016). So, once its position retrieved on the genome (HiC_scaffold_29: 11410869-11422911 + strand), we visualized its expression on IGV using the RNA-seq .bam files. Based on the higher expression of the 3’UTR exon, we could determine that CC2, CC7, CC8, CR2, RC6, RR10 are males, and CR3, CR6, RC4, RC7, RR6 and RR8 are females (Supplementary **Fig. S7**).

To assess the overall quality of the samples and to evaluate the potential impact of sex on gene expression in FAW, we performed a principal component analysis (PCA) and hierarchical clustering on RNA-seq data. These analyses did not show groupings based on sex (**Fig. S8)**. A thorough examination of the PCA plots revealed an absence of clear segregation between female and male samples across the entire genome, including the comprehensive chromosome set (**Fig. S8-A**), the HiC_scaffold_1 autosome (**Fig. S8-B**), and the sex chromosome, represented by HiC_scaffold_29 (**Fig. S8-C** and **D**). This lack of differentiation suggests that sex-biased gene expression exerts a negligible influence on the FAW larvae at the stage analyzed. Interestingly, the observed groupings were not determined by sex but were instead significantly driven by the effects of genotype. Specifically, on chromosome 29 (**Fig. S8-C** and **D**), the first principal component (PC1) effectively separated the hybrid genotypes, CR and RC, from the parental genotypes, CC (except CC7) and RR. Furthermore, the second principal component (PC2) distinguished individuals based on their maternal lineage, with those from the corn-strain motherline (CC and CR, again except CC7) forming one group and those from the rice-strain motherline (RR and RC) forming another. The inconsistency of CC7 sample is probably because there are two different haplotypes of the CC strain in our lab populations (Supplementary **Fig. S4**), which might cause a discrepancy in the expression level.

Nonetheless, these findings indicate that, irrespective of the potential for dosage compensation mechanisms in FAW, any sex-related expression bias is not evident in our PCA dataset, particularly concerning chromosome 29 (**Fig. S8-C** and **D**).

**Strain differentiation showed by trajectory clustering**

Two distinct clusters showed differential expression profiles between the FAW parental strains. As shown in **Figure S9A**, cluster 1 illustrated a pronounced over-expression of 69 DEGs in the rice strain (RR) relative to the corn strain (CC). The two reciprocal hybrid genotypes displayed intermediate expression patterns, suggesting a genetic interplay between these two strains.

Conversely, the cluster 8 in **Figure S9B** revealed a similar pattern of over-expression of 235 DEGs in the corn strain (CC) in comparison to the rice strain (RR). Again, the two hybrid genotypes were positioned at an intermediate expression level.

However, when we attempted to extract genes from these two clusters and performed ORA, we did not detect any enriched BP terms or KEGG pathways that surpassed the established statistical threshold. Upon revisiting the gene lists, we discovered that only about half of the genes were annotated in both instances, with these annotated genes detailed in **Supplementary Tables S3 and S4**, respectively. Among the over-expressed annotated genes specific to the RR strain (**Supplementary Table S3**), we identified genes associated with various biological processes such as metabolism, signal transduction, gene regulation, stress response, and detoxification. In the context of the CC strain, the over-expressed annotated genes are involved in metabolism, cellular signaling, gene regulation and detoxification functions (**Supplementary Table S4**).

**RR-effect genes by LRT**

In our examination of the gene list exhibiting overexpression in the RR strain of FAW, we identified a considerable number of genes that are linked to metabolic processes (**Supplementary Table S3**). These processes include cellular signaling, gene regulation, and detoxification mechanisms, which are fundamental to the physiological and adaptive responses of the organism. For example, trehalose metabolism in Lepidoptera has a role in energy production, development, and stress response (Tang *et al.*, 2018; Li *et al.*, 2024). Maltase enzymes are important for carbohydrate digestion and energy production (Carneiro *et al.*, 2004). Alcohol dehydrogenases are involved in ethanol metabolism and detoxification, showing implications for insecticide resistance and alcohol tolerance (Zhao *et al.*, 2020). stAR-related lipid transfer protein, involved in lipid metabolism, is crucial for cell membrane maintenance and hormone production (Niwa and Niwa, 2014), and is associated to the flight activity of long-distance flying insects (van der Horst *et al.*, 2002). The *SCOT-1* gene encodes the Succinyl-CoA:3-ketoacid coenzyme A transferase 1, a key enzyme acting in mitochondria in the utilization of ketone bodies, which are alternative energy substrates produced during the breakdown of fatty acids (Cantrell and Mohiuddin, 2024). This gene may be essential for the survival and adaptation of insects to their environments, particularly in conditions where carbohydrate resources are limited or during stages of high energy demand. Indeed, empirical studies have reported an elevated expression of mitochondrial genes within the RR strain compared to the CC strain, which may be correlated with the RR strain’s heightened energy demands for its mobility and a more extensive range of host infestation (Orsucci *et al.*, 2022). This enhanced mitochondrial activity may confer a metabolic advantage to the RR strain, supporting its active lifestyle and potentially contributing to its ecological success. Similar interpretation has also been proposed recently to explain the facilitated invasion of FAW due to novel mito-nuclear combination (Li *et al*., 2024).

**CC-effect genes by LRT**

In our analysis of the overexpressed gene list from the CC strain of FAW (**Supplementary Table S4**), we discovered a broader array of seemingly unrelated functions. On the one hand, we found many detoxification enzymes, such as Cytochrome P450 enzymes, which are involved in detoxification, hormone synthesis, and metabolism of xenobiotics. Glutathione S-transferase (GST) enzymes were also detected, which are also crucial for detoxification processes in insects. Carbonyl reductases are a class of enzymes that catalyze the reduction of carbonyl groups to hydroxyl groups using ATP or NADPH as a cofactor (Basri *et al.*, 2023). They play a crucial role in various biological processes, including the detoxification of xenobiotics, the synthesis of certain hormones and lipids, and the metabolism of drugs and other compounds (Carvalho *et al.*, 2013). The overexpression of detoxification-associated genes in CC vs. RR is consistent with our previous study comparting the two strains (Orsucci *et al.*, 2018). On the other hand, we found genes involved in different pathways of regulation. For example, caspases are crucial regulators of apoptosis, and play roles in development, immune responses, and response to environmental stressors in Lepidoptera (Romanelli *et al.*, 2016). Proteins containing SET and MYND domains are associated with chromatin remodeling and transcriptional regulation and play roles in development, gene expression, and stress responses in Lepidoptera (Spellmon *et al.*, 2015). We also found reverse transcriptases that are likely putative transposable elements annotated as host genes. This overexpression of transposable elements in CC versus RR has also been previously reported (Orsucci *et al.*, 2018).

Our analysis suggests that the primary genetic distinctions between the two strains of FAW are predominantly associated with cellular metabolic processes such as detoxification pathways. A subset of genes within these pathways has emerged as potential candidates that may underline these differences, including lipid metabolism-related genes, sorbitol dehydrogenase, members of GSTs, UGTs, and possibly carbonyl reductases. Despite these preliminary findings, the specific roles of these genes in the biology and physiology of FAW are yet to be elucidated.

###### Transcriptional differences between RR and CC

To further elucidate the differential gene expression patterns between the two FAW strains, we undertook a pairwise comparative analysis of CC versus RR. PCA of the RNA-seq data revealed a distinct clustering of samples based on FAW strain, with 27% explained variance in gene expression on PC2 (**Fig. S10A**). The differential expression analysis by DESeq2 (see **Methods**) detected 203 genes that were overexpressed in RR relative to CC, and conversely, 229 genes that were overexpressed in CC compared to RR (**Fig. S10B**).

###### Functional analysis of the DEGs between RR and CC

After retrieving all DEGs from both strains, we performed ORA for GO terms and KEGG pathways to identify enriched biological processes and metabolic pathways (**Fig. S11**). Although no enriched BP terms were detected for either strain, several KEGG pathways showed significant enrichment.

For the RR strain, when compared to the CC strain, enrichment was observed in the following KEGG pathways: “Pentose and glucuronate interconversions” (ko00040, adjusted *p*-value = 0.027), “Steroid biosynthesis” (ko00100, adjusted *p*-value = 0.029), “Metabolism of xenobiotics by cytochrome P450” (ko00980, adjusted *p*-value = 0.036), and “Glutathione metabolism” (ko00480, adjusted *p*-value = 0.036).

Interestingly, when contrasting the gene expression profiles of the CC strain versus the RR strain, we identified two KEGG pathways that were significantly enriched in both comparative analyses (albeit with distinct adjusted *p*-values), namely “Metabolism of xenobiotics by cytochrome P450” (ko00980, adjusted *p*-value = 0.0005) and “Pentose and glucuronate interconversions” (ko00040, adjusted *p*-value = 0.0078). This finding suggests a commonality in the metabolic processes between the strains, despite the differential gene regulation. The other enrichment in CC was noted in the following pathways: “Retinol metabolism” (ko00830, adjusted *p*-value = 0.0006), “Drug metabolism - cytochrome P450” (ko00982, adjusted *p*-value = 0.0013), “Ascorbate and aldarate metabolism” (ko00053, adjusted *p*-value = 0.0078), “Porphyrin metabolism” (ko00860, adjusted *p*-value = 0.0078), and “Drug metabolism - other enzymes” (ko00983, adjusted *p*-value = 0.042), which involves different metabolic mechanisms.

###### Candidate genes involved in the differential expression of FAW strains

Subsequently, we conducted a gene extraction from the four enriched KEGG pathways identified in RR compared to CC. A total of 11 genes were discovered (**Fig. S12**), including a few metabolic genes and some genes related to detoxification, among which there might be target genes that could explain the transcriptional differences between the two strains.

We then extracted the genes from the enriched KEGG pathways specific to the CC strain when compared to the RR strain of FAW, and we detected six genes that could potentially contribute to the differential gene expression observed within the CC strain (**Fig. S13**). These genes were implicated in metabolic processes (such as detoxification), including “Epidermal Retinol Dehydrogenase 2 Isoform X1”, “Glutathione S-Transferase 2-like”, “Carbonyl Reductase [NADPH] 3-like”, “UDP-Glucuronosyltransferase 33B13”, “UDP-Glucosyltransferase 2”, and “UDP-Glucuronosyltransferase 2B20-like isoform X1”.

### Supplementary Figures and Tables

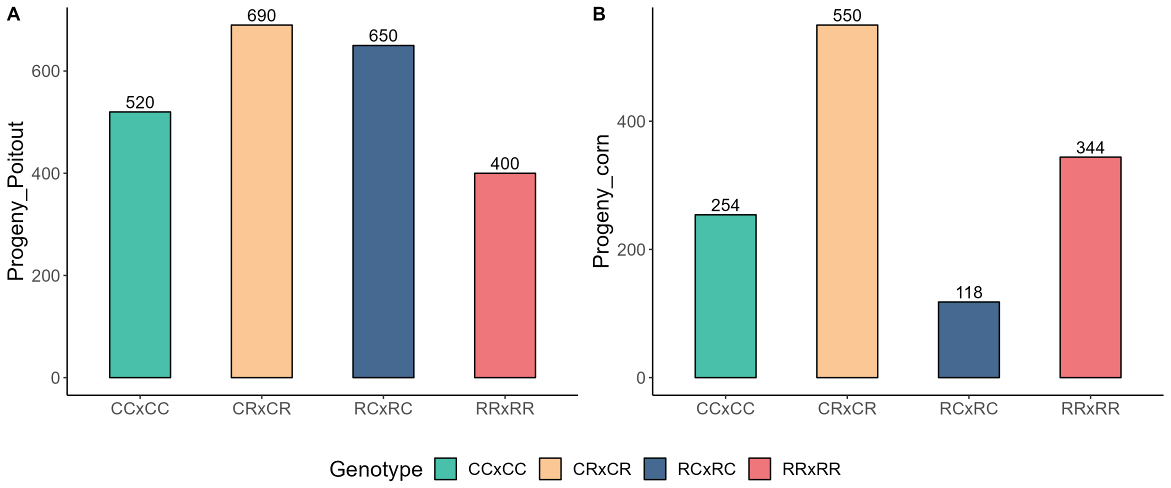

**Figure S1.** Different average offspring of the four FAW genotypes on both artificiat diet (A. Poitout) and corn plants (B). We did (five females × five males) for each genotype to increase the mating chance, and count the average progeny for each female (overall offspring number divided by five).

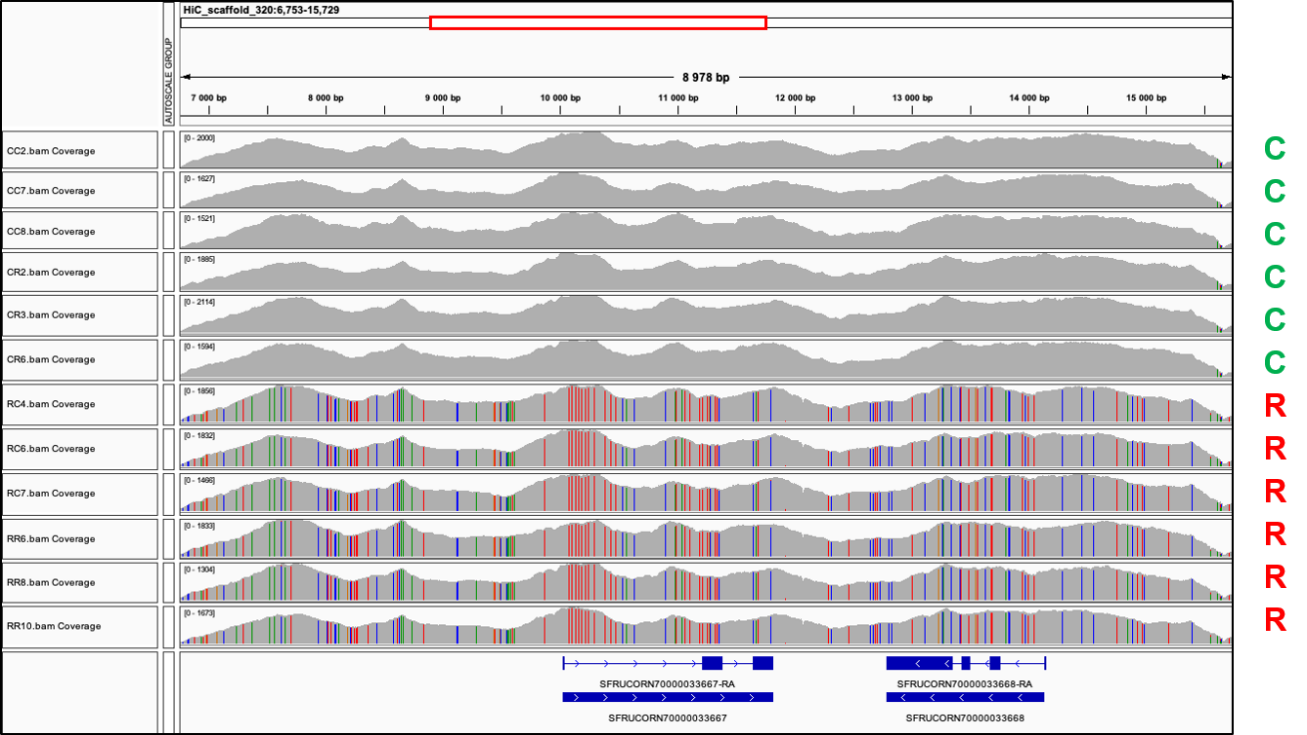

**Figure S2.** DNA re-sequencing of individuals of the four FAW genotypes visualized with IGV, showing the coverage of the 12 samples on the region 7000 bp – 15500 bp of the mitogenome (HiC_scaffold_320). The names of the 12 samples are indicated on the left, and the mitotypes are indicated on the right. Colored bars within the coverage tracks in gray represent SNPs. C mitotypes do not have any, compared to the corn-strain assembly, while R mitotypes show many SNPs.

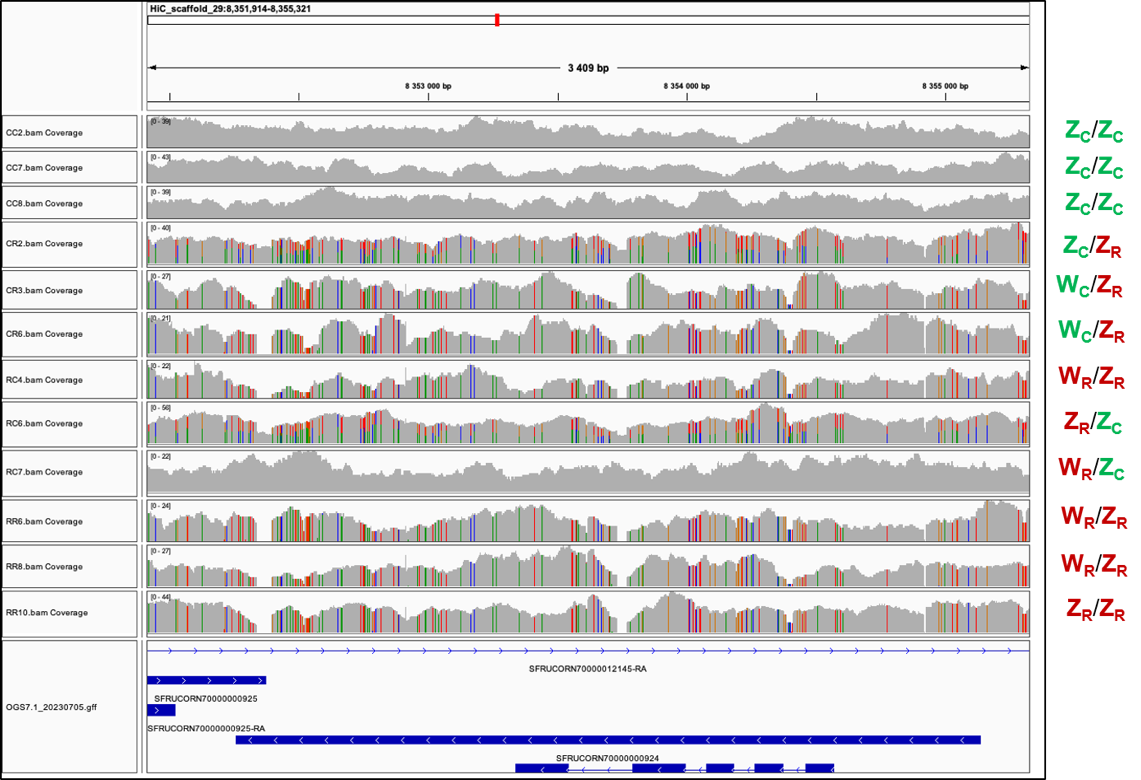

**Figure S3.** Re-sequencing of parental and hybrid individuals on the Z chromosome around the *Tpi* gene. Parental strains (CC and RR) exhibit only one genotype. Male hybrids (CR and RC) are heterozygous. Note: Screenshot from the IGV genome viewer represents the coverage of DNA-seq of the 12 samples on the region around the *Tpi* gene on the Z chromosome (HiC_scaffold_29: 8352000 bp – 8355500 bp). The names of the 12 samples are indicated on the left, and the genotypes are indicated on the right. Colored bars indicate SNPs and their proportion when heterozygous.

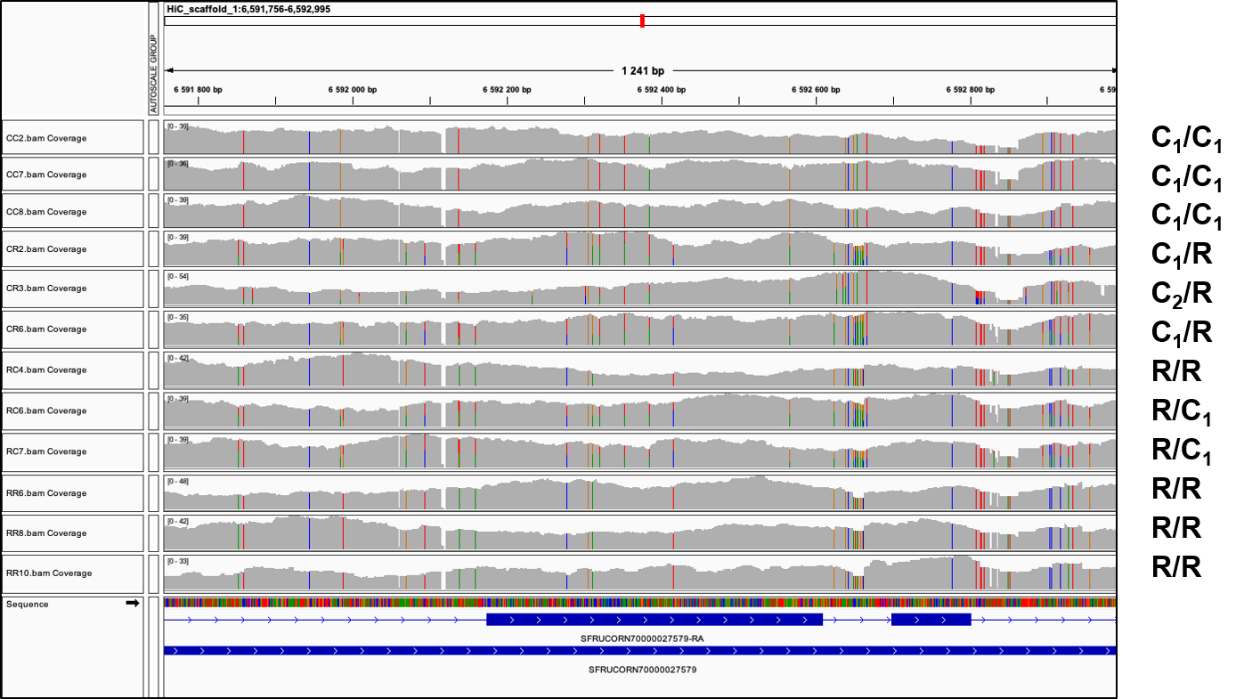

**Figure S4.** Screenshot from IGV showing the alignment of DNA-seq on an autosomal region (HiC_Scaffold_1: 6591800 bp – 6593000 bp). Colored bars indicate SNPs and their proportion when heterozygous. By checking the SNP composition for each individual in this region, CR3 shows a slightly different hybrid genotype than other CR individuals, due to a slightly different corn-strain haplotype, we named C2 in this figure. Only one haplotype can be observed from the rice strain in parental or hybrid individuals.

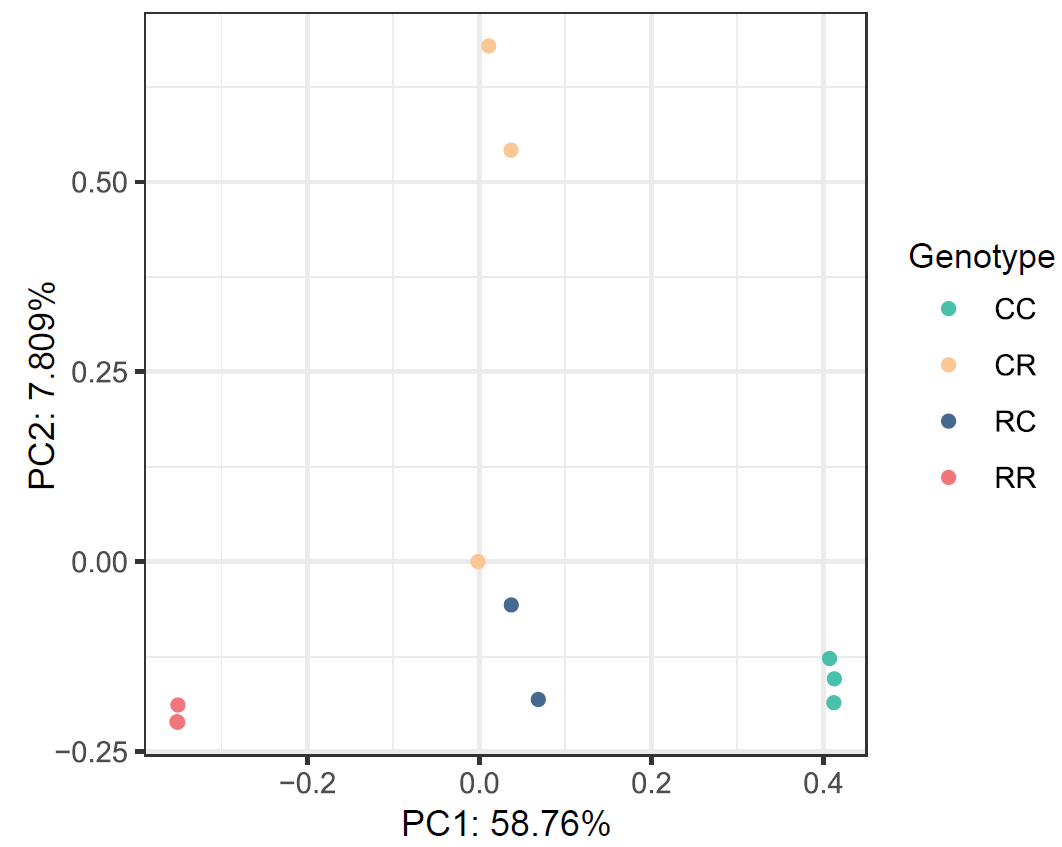

**Figure S5.** The result of principal component analysis (PCA) of whole genome sequencing on four FAW genotypes (represented by different colors).

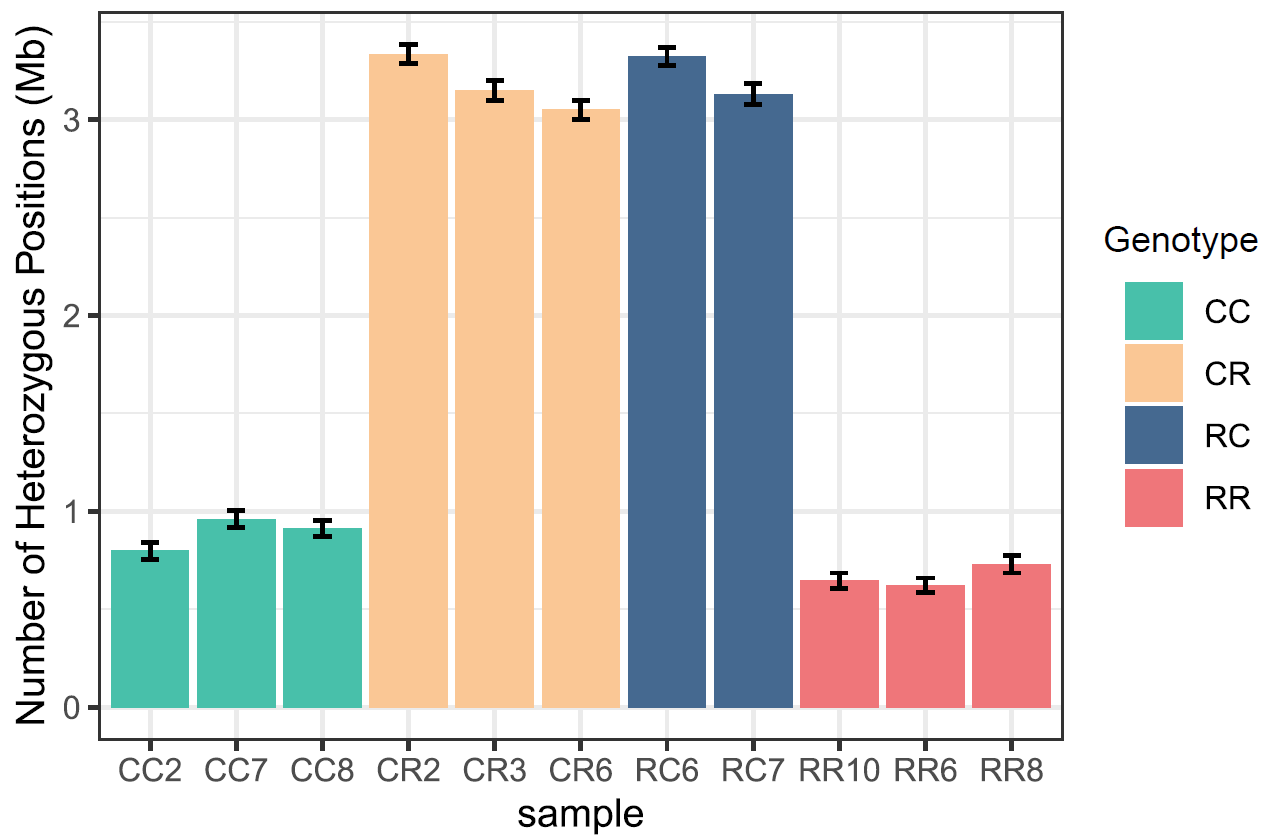

**Figure S6.** The number of heterozygous positions of four FAW genotypes (represented by different colors). The error bars indicate 95% confidence intervals calculated using non-parametric bootstrapping, resampling from 100 kb windows with 1,000 replications.

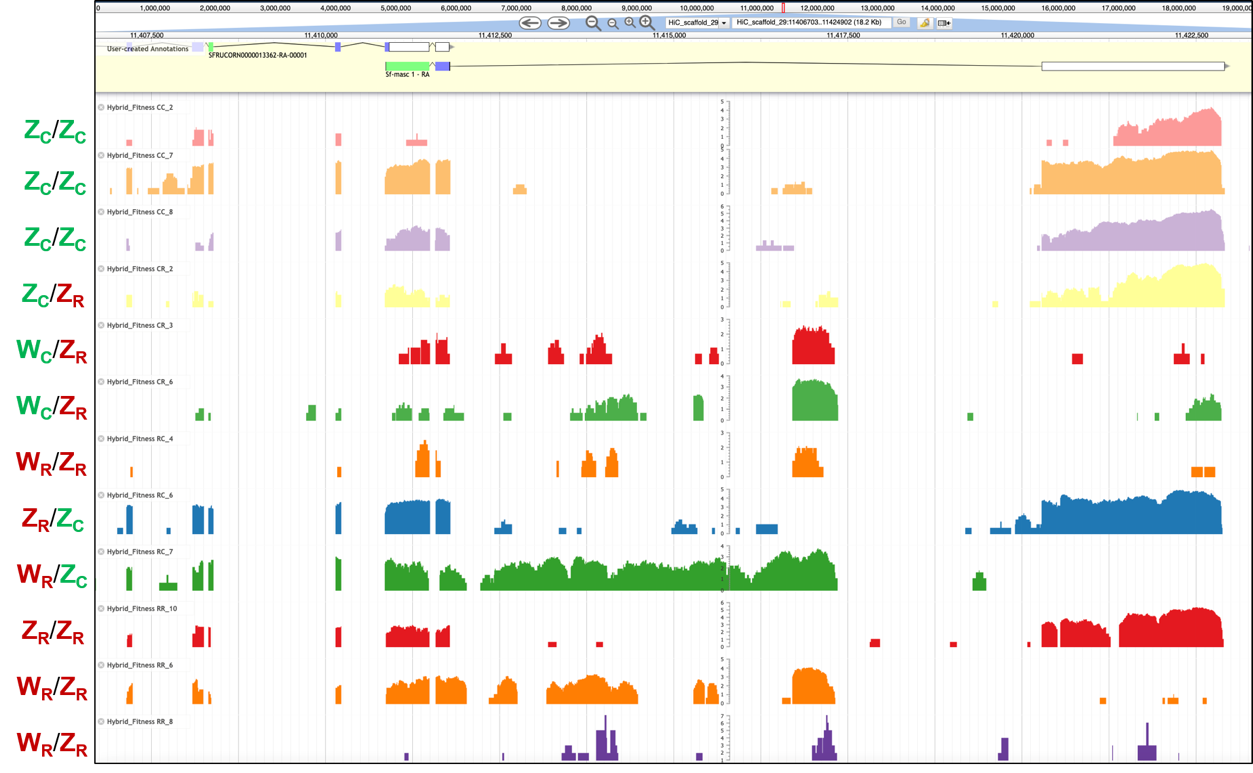

**Figure S7.** Expression of different genotypes on the 3’UTR exon. Note: Screenshot of the Apollo genome browser represents the coverage of RNA-seq of the 12 samples on the scaffold 29 between 11407500 bp and 11422500 bp. Sample names and genotypes are indicated on the left.

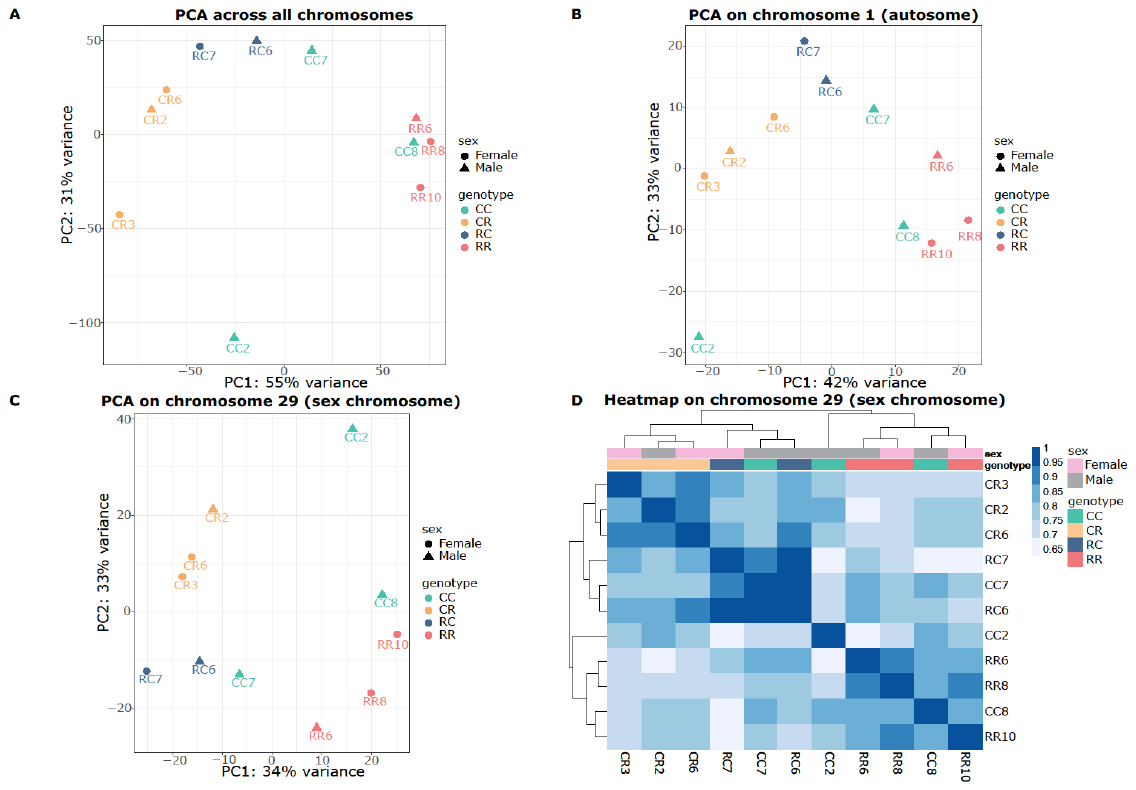

**Figure S8.** PCA plots and heatmap of RNA-seq dataset. The four genotypes are represented by different colors, and the sex are represented by different forms in PCA and by colors in the heatmap. Analysis of our genomic data revealed no significant clustering that aligned with sex, across the full spectrum of chromosomes within our genome background. This lack of sex-based segregation was consistent whether examining the genome as a whole (A), focusing on a specific autosome (B), or considering the sex chromosome (C & D) *per se*.

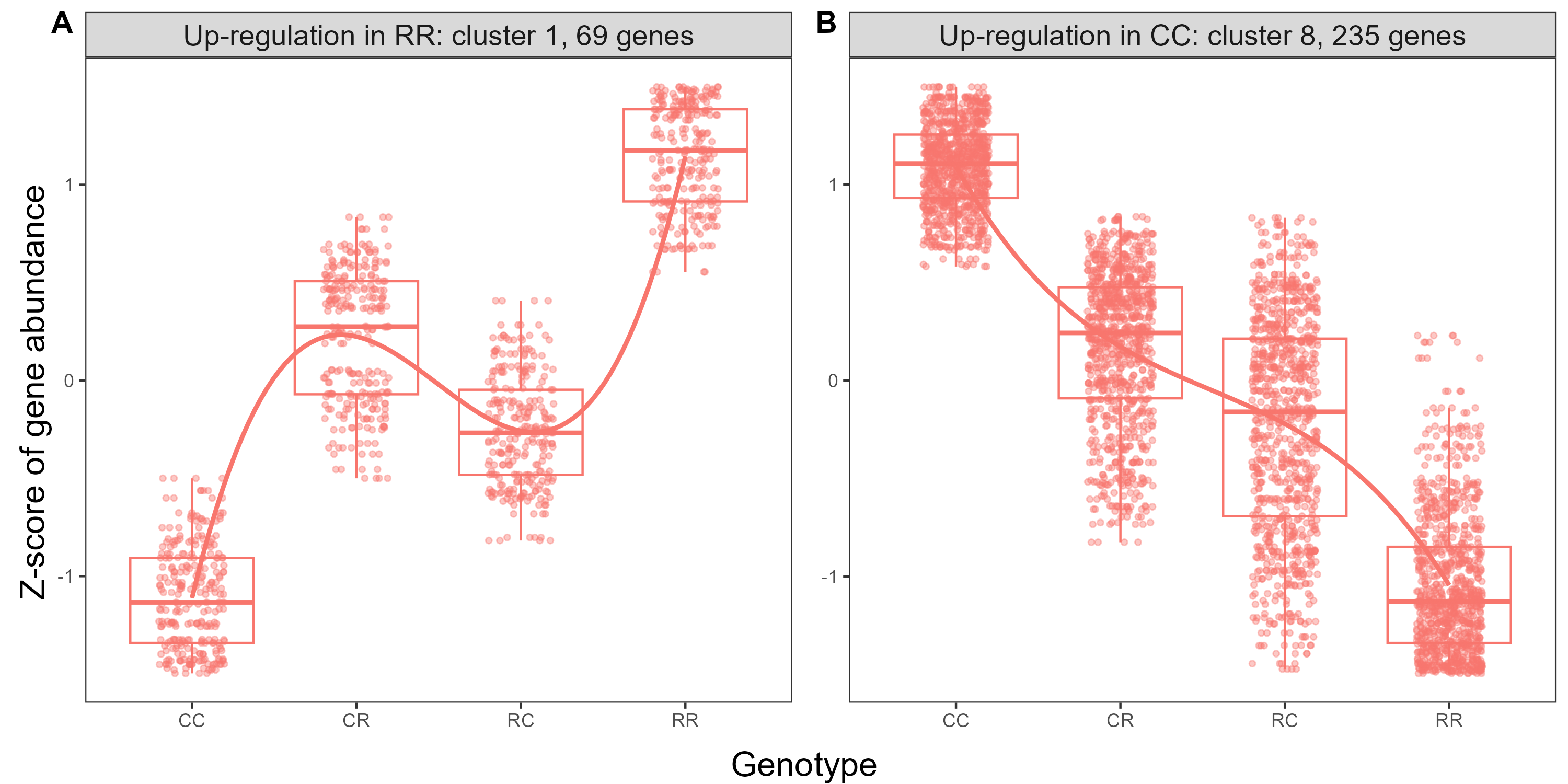

**Figure S9.** Selected clusters of DEGs showing typical differential expression patterns among the four FAW genotypes, especially between the two strains (CC and RR). (A). 69 up-regulated differentially expressed genes (DEGs) in RR compared CC; (B). 235 up-regulated DEGs in CC compared to RR. No enriched BP terms or KEGG pathways were identified within either of the two clusters under scrutiny. All gene expression clusters were given in Supplementary Figure **S15**.

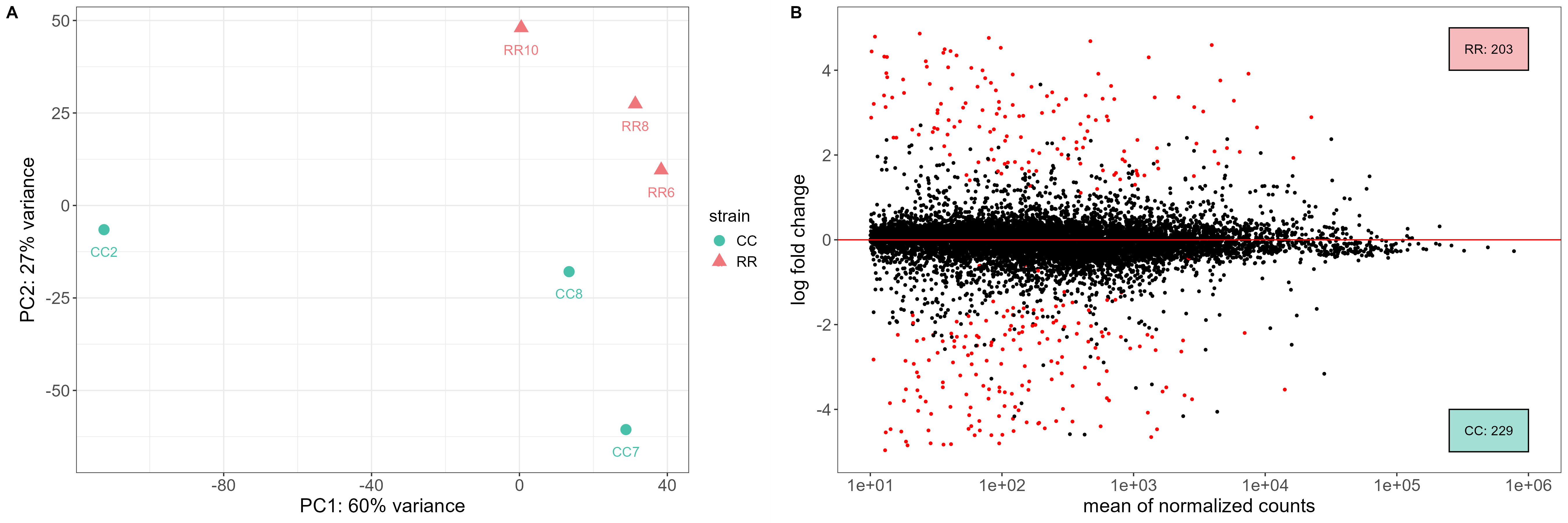

**Figure S10.** Transcriptional response of rice strain (RR) versus corn strain (CC). (A). Principal component analysis (PCA) on normalized RNA-seq reads for all the samples of CC and RR. (B). Multidimensional scaling plot (MA-plot) reporting the log2 fold changes between the RR and CC strains over the mean of normalized counts. Each dot in the MA-plot signifies an individual gene, with non-significant differential expression represented by black dots, and genes exhibiting significant differential expression marked by red dots. There are 203 up-regulated DEGs in RR compared to CC, and 229 up-regulated DEGs in CC compared to RR.

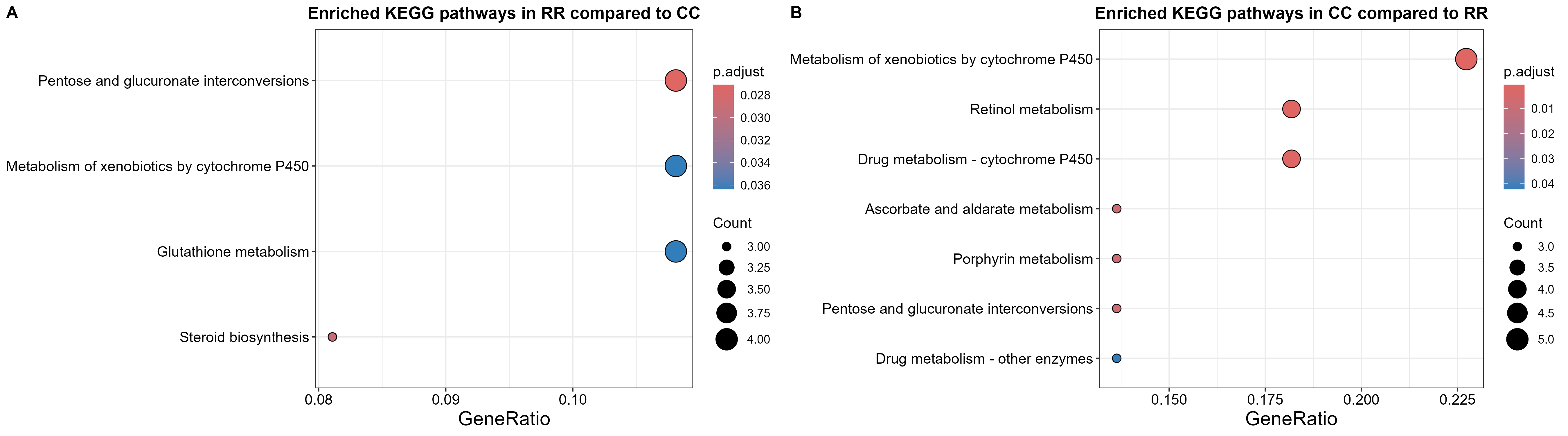

**Figure S11.** Enriched functional analysis of the DEGs for both RR strain and CC strain of FAW. (A). The enriched KEGG pathways in RR compared to CC; (B). The enriched KEGG pathways in CC compared to RR. Note: The size of black circles represents the number of genes implicated in each enriched KEGG pathway, and the adjusted *p*-value is represented by colors.

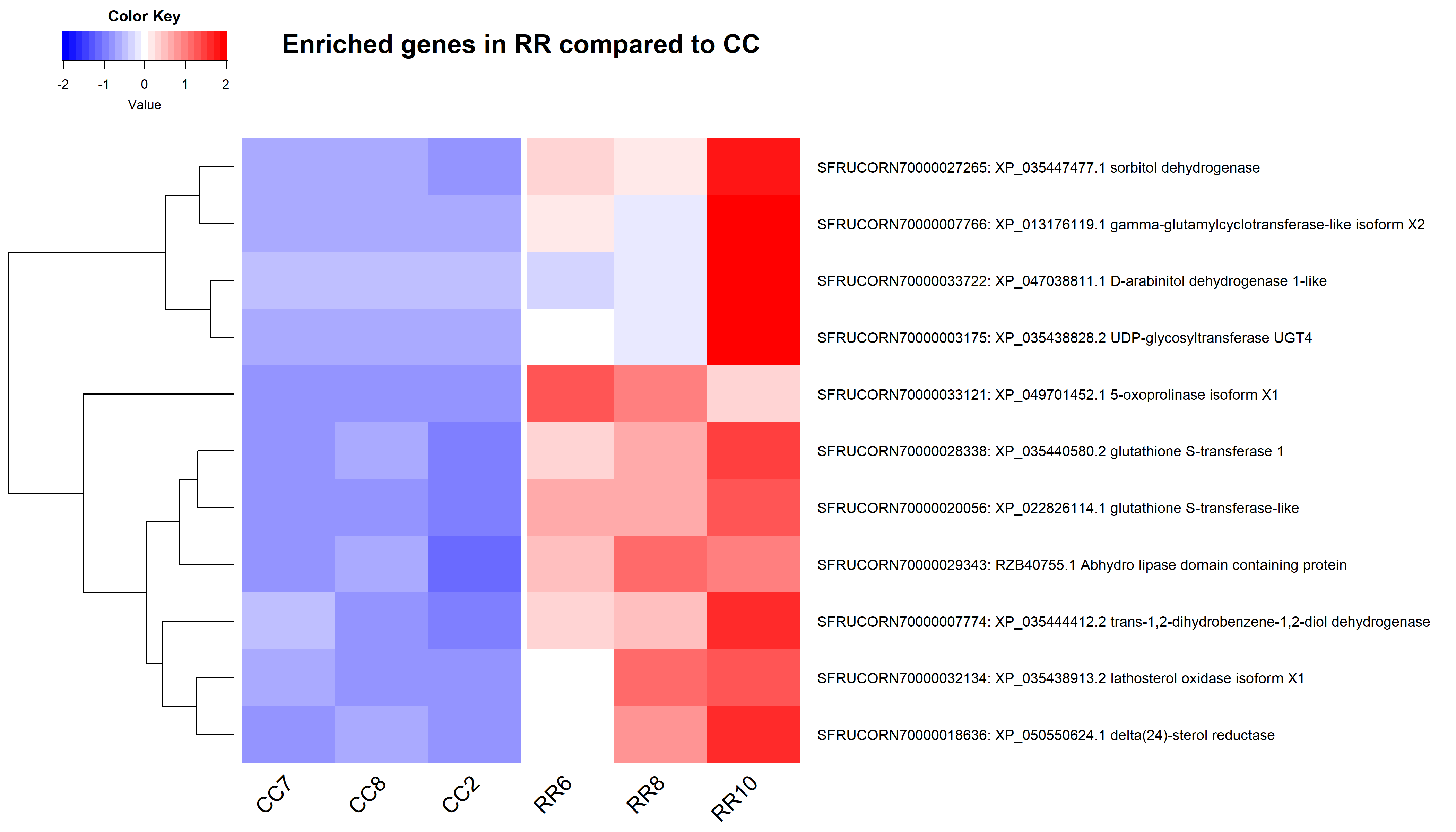

**Figure S12.** Heatmap showing the enriched genes that resulted from the differential expression of FAW rice strain compared to corn strain. Note: Scaled raw count values are represented by colors, name and function of each gene are indicated on the right.

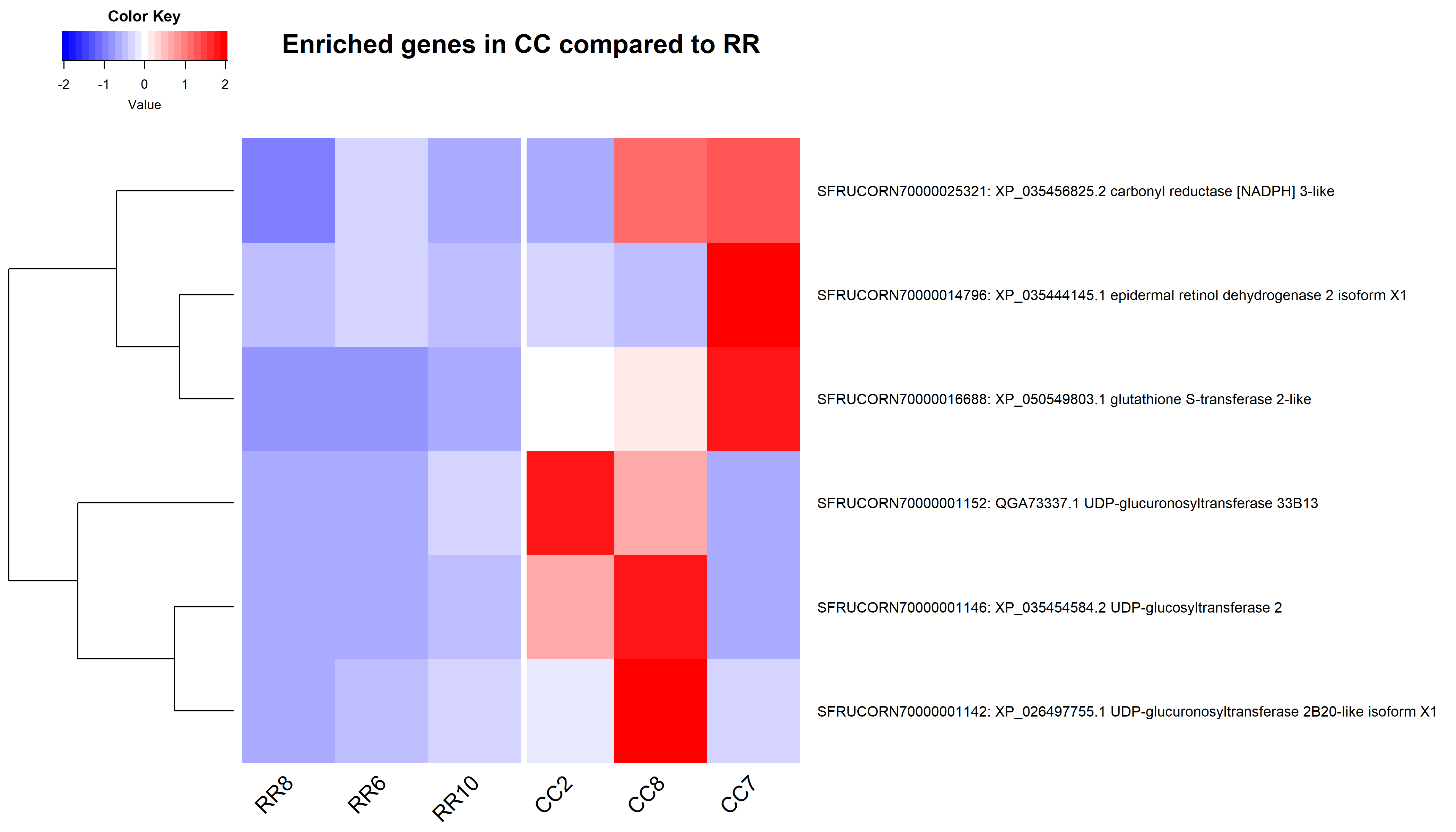

**Figure S13.** Heatmap showing the enriched genes that may result in the differential expression of FAW corn strain compared to rice strain. Note: Scaled raw count values are represented by colors, name and function of each gene are indicated on the right.

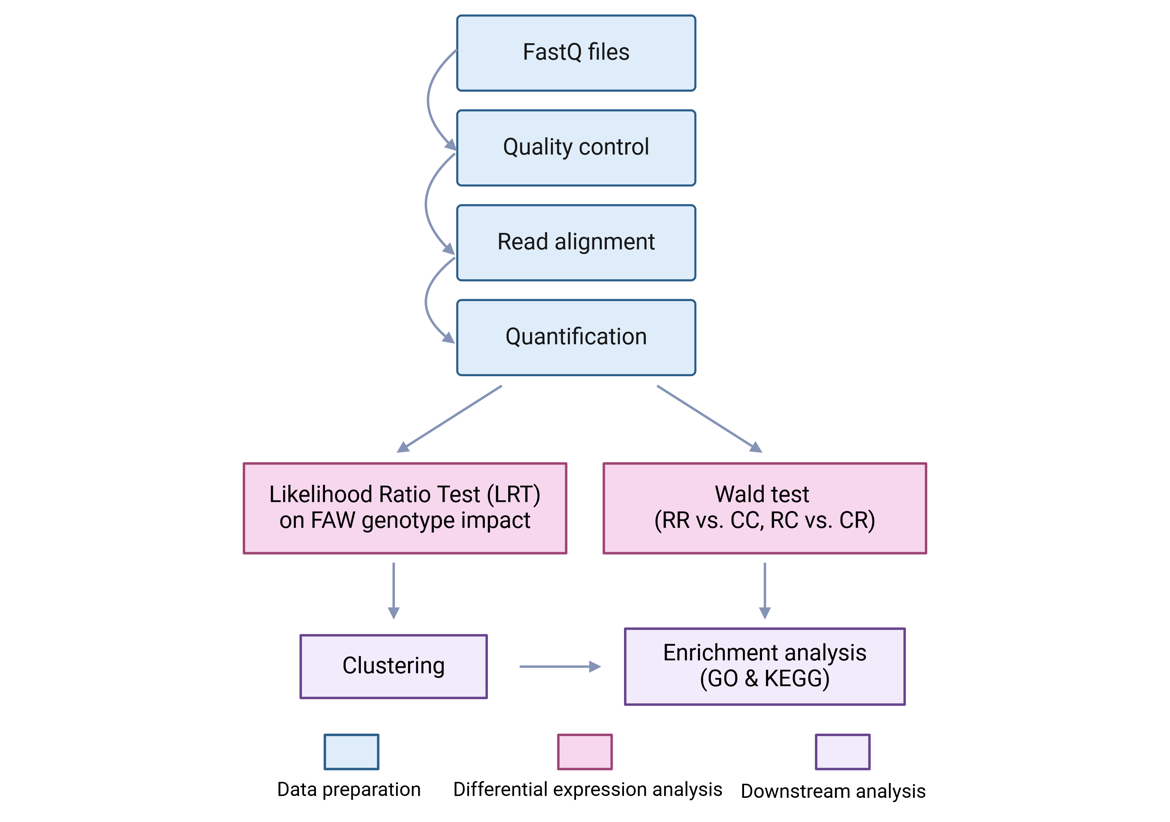

**Figure S14.** Overview of entire RNA-seq data analysis workflow. Adapted from (Greenig *et al.*, 2020). Created with (BioRender.com).

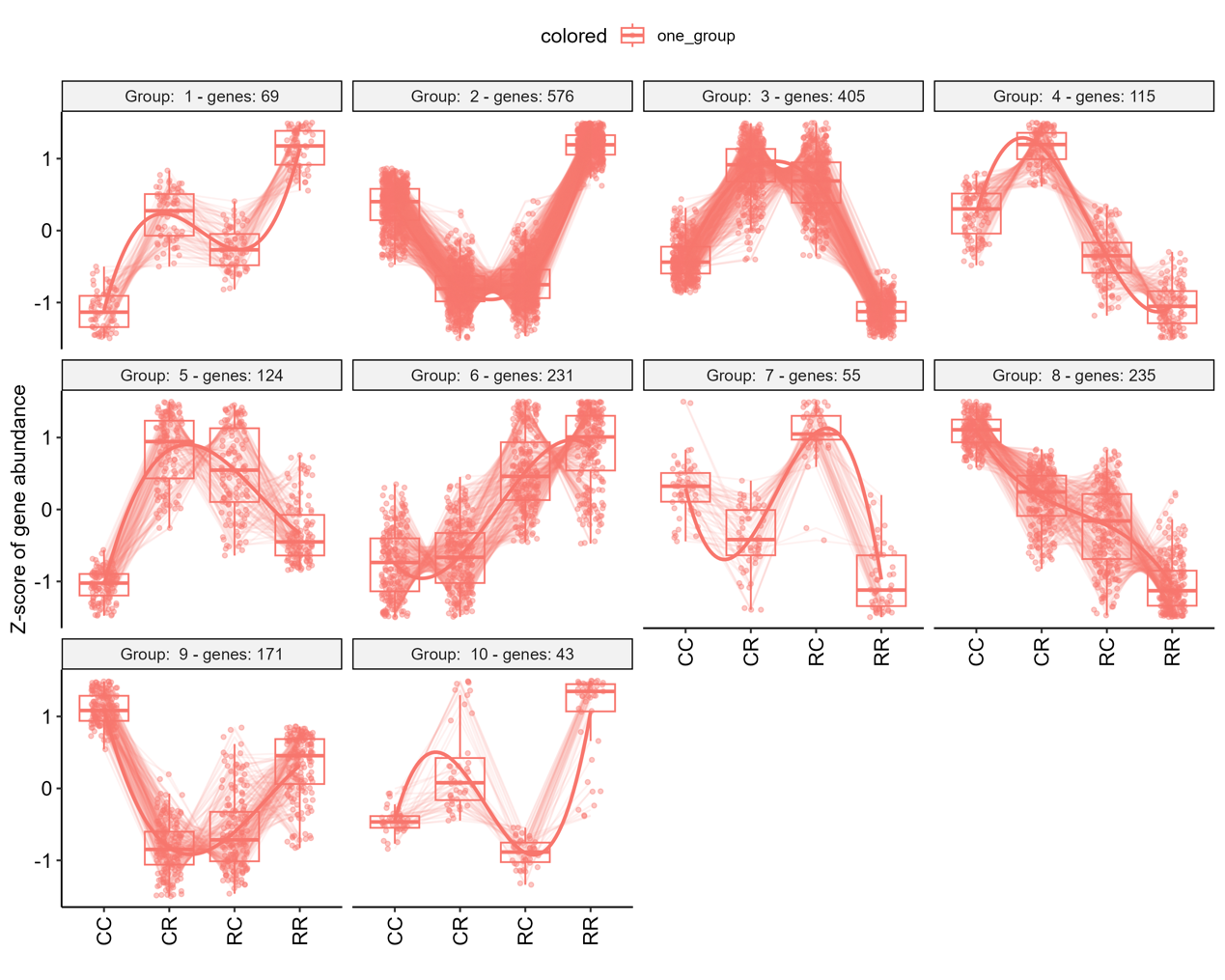

**Figure S15.** Entire gene expression clusters displaying the typical expression patterns of the resultant 10 clusters, as well as the scaled expression levels of each individual gene with DEGreport (v1.40.0, Pantano, 2024). Note: Each cluster is labeled with its number and the count of genes it contains, as indicated in the title of the corresponding scaled expression graph. The y-axis of the graph represents the genes plotted against their scaled expression values, measured by the Z-score of gene abundance, where values are centered to the mean and scaled to the standard deviation by each gene. The genotypes are indicated at the bottom. For each genotype, a box plot is included, with the median expression level marked by a horizontal line. Additionally, lines connect the mean expression levels across genotypes, illustrating the typical expression trajectory for each cluster.

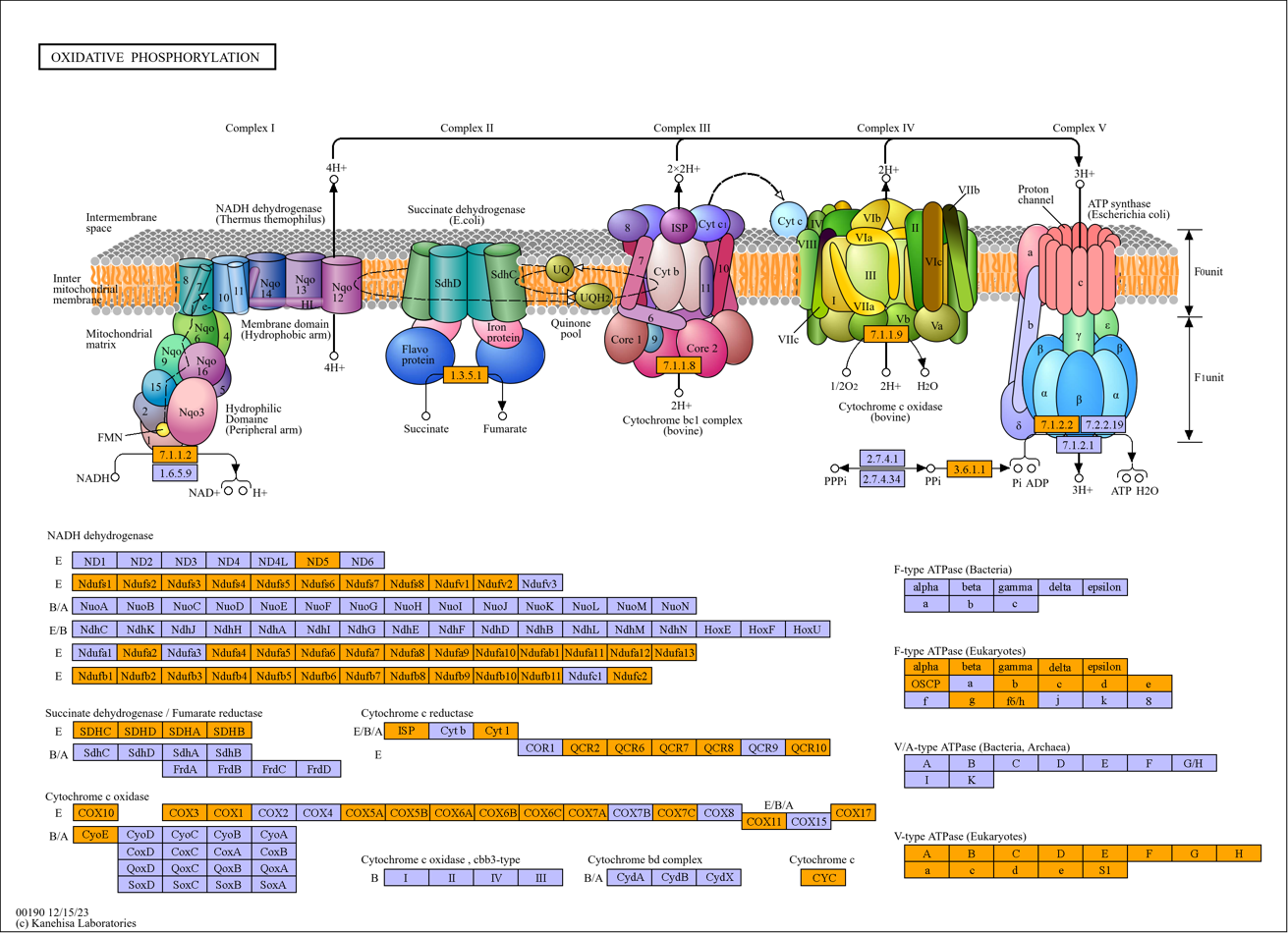

**Figure S16.** Illustration of the pathway map “oxidative phosphorylation” pathway (ko00190). For the genes that are present in our annotated genome in this pathway, they are marked in orange. Downloaded from (<https://www.genome.jp/entry/ko00190>).

**Table S1.** LRT_heterosis effect gene list

| **GeneID** | **Description** | **log2FoldChange** | **padj** |
| --- | --- | --- | --- |
| SFRUCORN70000002494 | XP_022832522.1peptidoglycan-recognition protein SC2-like isoform X3 | -0.16 | 2.35e-11 |
| SFRUCORN70000026492 | AFC87713.1Spod-11-tox b protein | -1.11 | 2.74e-07 |
| SFRUCORN70000024388 | XP_035454891.2NADPH oxidase 5 | -0.98 | 3.40e-07 |
| SFRUCORN70000001269 | XP_050557459.1protein spaetzle 5 | -1.78 | 1.41e-05 |
| SFRUCORN70000008350 | XP_050549581.1protein spaetzle 3 | -0.11 | 3.54e-05 |
| SFRUCORN70000006369 | AKJ54503.1cecropin B | -0.72 | 0.00022 |
| SFRUCORN70000023041 | XP_035429158.2neurotrophin 1 isoform X1 | -0.38 | 0.00060 |
| SFRUCORN70000028607 | XP_050559237.1dual oxidase isoform X1 | -0.91 | 0.0010 |
| SFRUCORN70000002209 | XP_035439049.1putative defense protein 3 | -4.96 | 0.0013 |
| SFRUCORN70000031411 | XP_050549682.1protein spaetzle 4 isoform X1 | -4.20 | 0.0015 |
| SFRUCORN70000005802 | XP_035438668.1protein spaetzle isoform X2 | -1.27 | 0.0032 |
| SFRUCORN70000001219 | AKJ54492.1attacin 2 | -2.90 | 0.036 |
| SFRUCORN70000011424 | KAG8111336.1hypothetical protein SFRUCORN_014347 | -2.88 | 0.00067 |
| SFRUCORN70000009798 | XP_035450645.1procathepsin L | -1.06 | 0.043 |
| SFRUCORN70000030419 | XP_035429207.2 catalase-like | -2.30 | 2.55e-12 |
| SFRUCORN70000030420 | XP_035429207.2 catalase-like | -6.38 | 2.08e-11 |
| SFRUCORN70000030418 | KAI5644031.1catalase domain-containing protein | -1.65 | 3.40e-11 |
| SFRUCORN70000002901 | OWR43125.1putative AlkB alkylation repair protein 2 | -0.95 | 9.50e-06 |
| SFRUCORN70000012649 | XP_037867691.1protein PFC0760c | -0.38 | 0.0056 |
| SFRUCORN70000007700 | XP_022820247.1TNF receptor-associated factor 4 isoform X2 | -0.90 | 0.013 |
| SFRUCORN70000014787 | XP_026746475.1mitogen-activated protein kinase kinase kinase 7-like isoform X1 | -0.42 | 0.020 |
| SFRUCORN70000002743 | XP_035439906.2transducin beta-like protein 2 isoform X2 | -0.82 | 0.020 |
| SFRUCORN70000010530 | XP_035451690.2alpha-crystallin A chain-like | -0.88 | 0.023 |
| SFRUCORN70000021295 | XP_035431816.1elongation of very long chain fatty acids protein 4 | -5.14 | 1.89e-08 |
| SFRUCORN70000029108 | XP_022816090.1elongation of very long chain fatty acids protein 7-like isoform X1 | -5.31 | 0.00035 |
| SFRUCORN70000033882 | XP_022834121.1stearoyl-CoA desaturase 5 | -0.96 | 0.0034 |
| SFRUCORN70000001489 | XP_035448342.2very-long-chain 3-oxoacyl-CoA reductase | -0.28 | 0.0079 |
| SFRUCORN70000023886 | XP_050557035.1elongation of very long chain fatty acids protein 7-like | -2.73 | 0.011 |

**Table S2.** LRT_maternal effect gene list

| **GeneID** | **Description** | **log2FoldChange** | **padj** |
| --- | --- | --- | --- |
| SFRUCORN70000033121 | XP_049701452.15-oxoprolinase isoform X1 | 8.12 | 2.31e-10 |
| SFRUCORN70000033695 | XP_035456298.2putative fatty acyl-CoA reductase CG5065 isoform X2 | 9.88 | 2.91e-06 |
| SFRUCORN70000020056 | XP_022826114.1glutathione S-transferase-like | 3.17 | 1.61e-05 |
| SFRUCORN70000001644 | XP_050552820.1luciferin 4-monooxygenase | 1.95 | 7.13e-05 |
| SFRUCORN70000019003 | XP_031768002.1 5-oxoprolinase-like | 6.51 | 0.00043 |
| SFRUCORN70000024755 | XP_035459295.1bifunctional 3'-phosphoadenosine 5'-phosphosulfate synthase isoform X1 | 2.12 | 0.0018 |
| SFRUCORN70000006351 | XP_035442077.2sulfotransferase 1B1-like | 3.20 | 0.0041 |
| SFRUCORN70000028081 | XP_050559192.1luciferin sulfotransferase-like | 2.29 | 0.0066 |
| SFRUCORN70000010961 | XP_035452115.1glutamate--cysteine ligase regulatory subunit | 1.65 | 0.021 |
| SFRUCORN70000017322 | XP_035437669.2persulfide dioxygenase ETHE1 | 1.40 | 0.023 |
| SFRUCORN70000032322 | XP_050559790.1 ATP-citrate synthase | 1.14 | 0.023 |
| SFRUCORN70000008555 | XP_050561167.1trimethyllysine dioxygenase | 3.65 | 0.0063 |

**Table S3.** LRT_RR up-regulated annotated genes

| **GeneID** | **Description** | **log2FoldChange** | **padj** |
| --- | --- | --- | --- |
| SFRUCORN70000019002 | XP_050552472.1tRNA (cytosine(34)-C(5))-methyltransferase | 10.28 | 4.05e-38 |
| SFRUCORN70000023334 | XP_050553878.1lactase/phlorizin hydrolase-like | 6.42 | 1.22e-19 |
| SFRUCORN70000033778 | XP_035441857.2facilitated trehalose transporter Tret1-like | 8.85 | 5.57e-15 |
| SFRUCORN70000023310 | XP_050553912.1spermatogenesis-defective protein 39 homolog | 6.92 | 3.22e-11 |
| SFRUCORN70000013859 | XP_050550725.1maltase A1-like | 7.16 | 2.84e-09 |
| SFRUCORN70000030737 | XP_050560116.1alcohol dehydrogenase 1-like | 8.06 | 8.47e-09 |
| SFRUCORN70000002074 | XP_021188627.1 U1 small nuclear ribonucleoprotein C | 3.45 | 2.38e-08 |
| SFRUCORN70000021306 | KAI5634393.1putative peptidase (DUF1758) domain-containing protein | 7.77 | 8.48e-08 |
| SFRUCORN70000008108 | XP_035441270.2putative carbonic anhydrase 5 | 5.06 | 2.87e-07 |
| SFRUCORN70000011486 | XP_050554898.1trypsin CFT-1-like | 4.97 | 3.92e-07 |
| SFRUCORN70000009895 | XP_022816178.1zinc finger CCHC domain-containing protein 3-like | 7.99 | 1.12e-06 |
| SFRUCORN70000020996 | XP_050559546.1stAR-related lipid transfer protein 7 | 2.08 | 3.76e-06 |
| SFRUCORN70000009061 | XP_035450397.2PI-PLC X domain-containing protein 3 | 3.18 | 5.15e-06 |
| SFRUCORN70000005814 | KAG8119067.1hypothetical protein SFRUCORN_021375 | 4.88 | 1.43e-05 |
| SFRUCORN70000024456 | XP_022819932.1 peripherin-2-like | 6.38 | 1.70e-05 |
| SFRUCORN70000010430 | XP_050555955.1ecdysone oxidase-like isoform X1 | 5.17 | 4.21e-05 |
| SFRUCORN70000013654 | XP_050559106.1myb/SANT-like DNA-binding domain-containing protein 4 isoform X2 | 3.85 | 6.74e-05 |
| SFRUCORN70000009303 | XP_035442404.2cytochrome P450 4c3-like | 5.23 | 1.99e-04 |
| SFRUCORN70000020984 | XP_035444820.2alanine aminotransferase 1-like | 1.55 | 2.52e-04 |
| SFRUCORN70000005806 | XP_035438562.1uncharacterized protein LOC118268256 | 2.18 | 4.99e-04 |
| SFRUCORN70000020981 | XP_035444669.1alanine aminotransferase 1 isoform X1 | 1.46 | 9.91e-04 |
| SFRUCORN70000024455 | XP_035429052.1succinyl-CoA:3-ketoacid coenzyme A transferase 1 | 1.77 | 1.53e-03 |
| SFRUCORN70000018307 | XP_035433913.1transcription initiation factor IIE subunit beta | 1.88 | 2.33e-03 |
| SFRUCORN70000019844 | XP_035443083.2cytochrome P450 6B5-like | 6.59 | 6.81e-03 |
| SFRUCORN70000017915 | XP_035447828.1alpha-tocopherol transfer protein-like isoform X2 | 2.96 | 8.87e-03 |
| SFRUCORN70000009722 | XP_050554271.1nose resistant to fluoxetine protein 6 | 6.37 | 8.98e-03 |
| SFRUCORN70000003972 | XP_050558165.1peroxisomal biogenesis factor 19-like isoform X2 | 2.42 | 1.55e-02 |
| SFRUCORN70000017390 | XP_035440548.2valacyclovir hydrolase | 1.85 | 1.60e-02 |
| SFRUCORN70000020413 | XP_035444951.2L-lactate dehydrogenase isoform X1 | 2.05 | 1.77e-02 |
| SFRUCORN70000010479 | KAG8114460.1hypothetical protein SFRUCORN_015749 | 2.11 | 2.03e-02 |
| SFRUCORN70000007065 | XP_022828706.1cysteine-rich PDZ-binding protein-like | 4.06 | 3.29e-02 |
| SFRUCORN70000006201 | XP_050550949.1myb/SANT-like DNA-binding domain-containing protein 4 | 3.77 | 3.57e-02 |
| SFRUCORN70000002184 | XP_035438917.2corticotropin-releasing factor-binding protein | 2.40 | 4.50e-02 |
| SFRUCORN70000022876 | XP_035450049.2peroxisomal catalase 1-like | 3.03 | 4.73e-02 |

**Table S4.** LRT_CC up-regulated annotated genes

| **GeneID** | **Description** | **log2FoldChange** | **padj** |
| --- | --- | --- | --- |
| SFRUCORN70000023673 | XP_050560898.1probable cytochrome P450 9f2 | -5.44 | 1.05e-09 |
| SFRUCORN70000022203 | KAG8102657.1hypothetical protein SFRUCORN_019431 | -2.86 | 1.14e-09 |
| SFRUCORN70000033926 | XP_037295504.1caspase Dronc-like isoform X1 | -10.63 | 2.80e-09 |
| SFRUCORN70000009773 | XP_050554246.1PRKCA-binding protein isoform X1 | -5.91 | 2.97e-09 |
| SFRUCORN70000023994 | WP_047927799.1reverse transcriptase domain-containing protein | -8.43 | 3.18e-09 |
| SFRUCORN70000016690 | XP_035444263.2SET and MYND domain-containing protein 4 | -5.70 | 4.33e-09 |
| SFRUCORN70000009396 | XP_037295504.1caspase Dronc-like isoform X1 | -9.31 | 4.44e-09 |
| SFRUCORN70000026251 | XP_037295504.1caspase Dronc-like isoform X1 | -9.35 | 4.50e-09 |
| SFRUCORN70000024807 | XP_026725659.1caltractin-like isoform X1 | -4.48 | 9.44e-09 |
| SFRUCORN70000023739 | XP_035440836.1transmembrane protease serine 9 | -9.49 | 1.78e-08 |
| SFRUCORN70000010154 | XP_035432872.2autophagy protein 12-like | -2.61 | 3.94e-08 |
| SFRUCORN70000003724 | KAG8108368.1hypothetical protein SFRUCORN_000250 | -8.03 | 7.75e-08 |
| SFRUCORN70000028084 | XP_035436318.2sulfotransferase 1C4 isoform X1 | -3.77 | 7.78e-08 |
| SFRUCORN70000023679 | AID55431.1cytochrome P450 9A60 | -2.45 | 3.16e-07 |
| SFRUCORN70000011482 | XP_050554893.1trypsin CFT-1-like | -5.50 | 1.55e-06 |
| SFRUCORN70000017658 | XP_050558594.1sodium/nucleoside cotransporter 2-like | -2.99 | 2.86e-06 |
| SFRUCORN70000006782 | XP_022835360.1lipase member I-like | -7.23 | 3.71e-06 |
| SFRUCORN70000023049 | XP_035443518.2 putative nuclease HARBI1 | -4.99 | 9.57e-06 |
| SFRUCORN70000018590 | XP_047020359.1chorion peroxidase-like | -7.21 | 3.54e-05 |
| SFRUCORN70000018656 | XP_035435233.2nose resistant to fluoxetine protein 6-like | -6.56 | 4.71e-05 |
| SFRUCORN70000014518 | KAG8105059.1hypothetical protein SFRUCORN_002012 | -5.02 | 6.09e-05 |
| SFRUCORN70000024458 | XP_004931922.1protein PET117 homolog | -2.34 | 8.22e-05 |
| SFRUCORN70000003606 | XP_022827731.1antichymotrypsin-2-like isoform X1 | -3.19 | 9.38e-05 |
| SFRUCORN70000005337 | XP_035439410.1ATP synthase subunit gamma | -5.08 | 1.05e-04 |
| SFRUCORN70000025321 | XP_035456825.2carbonyl reductase [NADPH] 3-like | -2.41 | 2.65e-04 |
| SFRUCORN70000030757 | XP_022827656.1probable 28S ribosomal protein S25 | -2.62 | 3.33e-04 |
| SFRUCORN70000010660 | XP_031763831.1heat shock transcription factor | -6.32 | 4.17e-04 |
| SFRUCORN70000003421 | XP_035447696.2alcohol dehydrogenase-like | -2.89 | 5.54e-04 |
| SFRUCORN70000019525 | XP_050556908.1solute carrier family 2 | -2.59 | 6.66e-04 |
| SFRUCORN70000003728 | XP_035434647.2glutamyl-tRNA(Gln) amidotransferase subunit B | -2.64 | 7.08e-04 |
| SFRUCORN70000015816 | CAH0689981.1unnamed protein product | -2.59 | 7.10e-04 |
| SFRUCORN70000007504 | KAG8120782.1hypothetical protein SFRUCORN_004012 | -2.43 | 9.11e-04 |
| SFRUCORN70000026539 | XP_050559734.1GILT-like protein 1 isoform X1 | -2.53 | 1.11e-03 |
| SFRUCORN70000011633 | KAG8101251.1hypothetical protein SFRUCORN_018032 | -7.40 | 1.20e-03 |
| SFRUCORN70000014162 | XP_050558361.1protein KTI12 homolog | -2.42 | 1.52e-03 |
| SFRUCORN70000021855 | XP_035437366.2gastrula zinc finger protein XlCGF57.1 | -2.64 | 1.57e-03 |
| SFRUCORN70000016192 | XP_035453132.1uncharacterized protein LOC118278152 isoform X1 | -1.28 | 1.72e-03 |
| SFRUCORN70000022112 | XP_022817897.1ribonuclease P protein subunit p21 | -1.70 | 1.77e-03 |
| SFRUCORN70000022461 | XP_035442176.1charged multivesicular body protein 3 | -2.93 | 2.06e-03 |
| SFRUCORN70000020436 | XP_035441839.1ER membrane protein complex subunit 6 | -1.68 | 2.13e-03 |
| SFRUCORN70000030002 | KAI5638843.1baculovirus F protein domain-containing protein | -7.20 | 3.13e-03 |
| SFRUCORN70000024070 | XP_035431876.2HORMA domain-containing protein 1-like | -6.83 | 3.52e-03 |
| SFRUCORN70000009579 | XP_050561979.1putative fatty acyl-CoA reductase CG5065 | -4.26 | 3.52e-03 |
| SFRUCORN70000007516 | XP_035445023.1ER membrane protein complex subunit 8/9 homolog | -1.27 | 3.92e-03 |
| SFRUCORN70000001284 | XP_022816700.1uracil phosphoribosyltransferase homolog | -1.46 | 4.19e-03 |
| SFRUCORN70000024094 | KAG8119635.1hypothetical protein SFRUCORN_008390 | -5.51 | 4.26e-03 |
| SFRUCORN70000004168 | XP_026725306.1chaoptin-like isoform X2 | -3.98 | 4.51e-03 |
| SFRUCORN70000029435 | KAG8114219.1hypothetical protein SFRUCORN_012224 | -1.43 | 4.70e-03 |
| SFRUCORN70000029182 | XP_022815155.1farnesyl pyrophosphate synthase 2-like | -4.86 | 5.22e-03 |
| SFRUCORN70000007788 | KPJ08750.15'-nucleotidase domain-containing protein 3 | -1.38 | 5.23e-03 |
| SFRUCORN70000016421 | KAG8102932.1hypothetical protein SFRUCORN_016435 | -2.97 | 5.51e-03 |
| SFRUCORN70000021310 | XP_035431846.2sucrose-6-phosphate hydrolase-like | -5.00 | 5.60e-03 |
| SFRUCORN70000031351 | XP_035452227.1guanine nucleotide exchange factor MSS4 homolog | -1.79 | 5.78e-03 |
| SFRUCORN70000017451 | XP_050554408.1S-methyl-5'-thioadenosine phosphorylase isoform X1 | -1.72 | 6.34e-03 |
| SFRUCORN70000022039 | XP_035456350.2arginine-hydroxylase NDUFAF5 | -1.51 | 6.59e-03 |
| SFRUCORN70000032692 | XP_050553298.11-acyl-sn-glycerol-3-phosphate acyltransferase beta isoform X2 | -3.81 | 6.92e-03 |
| SFRUCORN70000019232 | KAG8111884.1hypothetical protein SFRUCORN_008826 | -2.32 | 7.05e-03 |
| SFRUCORN70000010830 | XP_050551567.1phosphatidylglycerophosphatase and protein-tyrosine phosphatase 1 | -1.77 | 7.07e-03 |
| SFRUCORN70000021753 | XP_035443325.2GDP-D-glucose phosphorylase 1 | -1.29 | 7.16e-03 |
| SFRUCORN70000006582 | XP_035433136.2deoxyhypusine hydroxylase | -1.84 | 7.39e-03 |
| SFRUCORN70000015039 | XP_035430601.1adenylate kinase 8 | -3.82 | 7.61e-03 |
| SFRUCORN70000032835 | XP_035431846.2sucrose-6-phosphate hydrolase-like | -4.82 | 9.50e-03 |
| SFRUCORN70000018921 | XP_038219659.1inositol phosphorylceramide glucuronosyltransferase 1-like | -2.49 | 1.01e-02 |
| SFRUCORN70000008964 | XP_035448270.1potassium channel subfamily K member 15 | -1.57 | 1.10e-02 |
| SFRUCORN70000005521 | XP_035449367.2RAB6A-GEF complex partner protein 2 | -1.23 | 1.18e-02 |
| SFRUCORN70000016508 | XP_035452629.1cyclin-dependent kinases regulatory subunit-like | -1.56 | 1.21e-02 |
| SFRUCORN70000001640 | XP_050552826.1cytochrome P450 4C1 | -6.29 | 1.24e-02 |
| SFRUCORN70000001224 | XP_035454848.2nuclear pore glycoprotein p62-like isoform X1 | -1.29 | 1.29e-02 |
| SFRUCORN70000022294 | QGA73351.1DnaJ-like protein 60 | -1.86 | 1.34e-02 |
| SFRUCORN70000002708 | XP_035440046.2tRNA (cytosine(72)-C(5))-methyltransferase NSUN6 | -2.34 | 1.37e-02 |
| SFRUCORN70000011866 | XP_050552761.1DNA cross-link repair 1A protein isoform X1 | -1.97 | 1.37e-02 |
| SFRUCORN70000033907 | ASN63931.1glutathione S-transferase delta 2 | -1.56 | 1.38e-02 |
| SFRUCORN70000022681 | XP_026326846.1probable protein kinase DDB_G0291133 | -8.78 | 1.53e-02 |
| SFRUCORN70000024632 | XP_035449558.2DNA polymerase epsilon catalytic subunit 1 | -1.31 | 1.60e-02 |
| SFRUCORN70000015375 | XP_035452051.2dysbindin protein homolog | -1.38 | 1.87e-02 |
| SFRUCORN70000029041 | XP_022826095.1lachesin-like isoform X2 | -3.21 | 1.89e-02 |
| SFRUCORN70000024457 | XP_035429056.139S ribosomal protein L20 | -1.08 | 2.05e-02 |
| SFRUCORN70000015410 | XP_050552394.1sodium/hydrogen exchanger 10 | -1.68 | 2.07e-02 |
| SFRUCORN70000012542 | XP_050559866.1titin homolog isoform X2 | -2.96 | 2.13e-02 |
| SFRUCORN70000024499 | XP_050558941.1S1 RNA-binding domain-containing protein 1 isoform X2 | -1.27 | 2.13e-02 |
| SFRUCORN70000012192 | XP_050561706.1RNA-binding protein 42 isoform X2 | -1.33 | 2.14e-02 |
| SFRUCORN70000015742 | KAF9801824.1hypothetical protein SFRURICE_019703 | -2.85 | 2.16e-02 |
| SFRUCORN70000018417 | XP_035452769.1FUN14 domain-containing protein 1 isoform X1 | -1.62 | 2.19e-02 |
| SFRUCORN70000010585 | XP_035453491.2bifunctional arginine demethylase and lysyl-hydroxylase PSR | -1.17 | 2.24e-02 |
| SFRUCORN70000017960 | XP_035448211.1DNA-directed RNA polymerase III subunit RPC10 | -1.99 | 2.40e-02 |
| SFRUCORN70000027839 | XP_028178687.1activity-regulated cytoskeleton associated protein 2-like | -4.12 | 2.46e-02 |
| SFRUCORN70000015634 | XP_035435775.1sodium-coupled monocarboxylate transporter 2 | -3.56 | 2.46e-02 |
| SFRUCORN70000016333 | XP_022816281.1nuclear transcription factor Y subunit B-4 isoform X1 | -1.15 | 2.55e-02 |
| SFRUCORN70000008796 | XP_035431795.2alpha-tocopherol transfer protein-like | -3.07 | 2.70e-02 |
| SFRUCORN70000012089 | UYA18667.1 odorant binding protein 6 | -2.07 | 2.85e-02 |
| SFRUCORN70000020719 | XP_035432672.2sulfotransferase 1 family member D1-like | -2.29 | 2.99e-02 |
| SFRUCORN70000009383 | XP_050552154.1lactase/phlorizin hydrolase-like | -2.44 | 3.05e-02 |
| SFRUCORN70000027637 | XP_050556148.1tRNA (adenine(58)-N(1))-methyltransferase catalytic subunit TRMT61A | -2.16 | 3.11e-02 |
| SFRUCORN70000030545 | XP_050556707.1exopolyphosphatase PRUNE1-like isoform X1 | -1.64 | 3.17e-02 |
| SFRUCORN70000013389 | XP_035439575.1uncharacterized protein LOC118268891 | -4.44 | 3.23e-02 |
| SFRUCORN70000011759 | XP_050559625.1probable WRKY transcription factor protein 1 isoform X2 | -2.91 | 3.35e-02 |
| SFRUCORN70000015969 | XP_022822184.1BLOC-1-related complex subunit 8 homolog | -1.38 | 3.43e-02 |
| SFRUCORN70000031529 | XP_026326846.1probable protein kinase DDB_G0291133 | -8.06 | 3.83e-02 |
| SFRUCORN70000020922 | XP_035442639.2probable asparagine--tRNA ligase | -1.53 | 3.93e-02 |
| SFRUCORN70000029924 | XP_022816147.1E3 SUMO-protein ligase NSE2-like | -1.91 | 4.11e-02 |
| SFRUCORN70000003502 | XP_035430007.139S ribosomal protein L51 | -1.80 | 4.12e-02 |
| SFRUCORN70000018383 | XP_035442109.228S ribosomal protein S18c | -2.00 | 4.22e-02 |
| SFRUCORN70000005592 | XP_035449277.2NPC intracellular cholesterol transporter 2-like | -2.34 | 4.25e-02 |
| SFRUCORN70000005682 | XP_050562390.1Fanconi anemia group M protein | -2.26 | 4.26e-02 |
| SFRUCORN70000012360 | XP_050563790.1microfibrillar-associated protein 1 | -1.40 | 4.31e-02 |
| SFRUCORN70000016913 | XP_022837887.1cuticle protein 10.9-like | -2.69 | 4.33e-02 |
| SFRUCORN70000003505 | XP_035429576.2ubiquinone biosynthesis protein COQ4 homolog | -1.21 | 4.35e-02 |
| SFRUCORN70000003061 | XP_035436723.2DNA repair protein RAD51 homolog 4 | -2.23 | 4.48e-02 |
| SFRUCORN70000024005 | XP_035433513.128S ribosomal protein S21 | -1.52 | 4.60e-02 |
| SFRUCORN70000023712 | XP_022814053.1GTP-binding protein 128up | -1.17 | 4.68e-02 |
| SFRUCORN70000008481 | TKX27959.1cuticular protein RR-2 | -2.43 | 4.71e-02 |
| SFRUCORN70000021471 | XP_035433970.2probable ATP-dependent RNA helicase DDX28 | -1.88 | 4.73e-02 |
| SFRUCORN70000002847 | XP_050559771.1general transcription factor IIH subunit 2 | -1.11 | 4.78e-02 |
| SFRUCORN70000001379 | XP_035454812.2probable 28S ribosomal protein S16 | -1.77 | 4.81e-02 |
| SFRUCORN70000027066 | KAG8102216.1hypothetical protein SFRUCORN_006580 | -6.15 | 4.82e-02 |
| SFRUCORN70000017282 | XP_035440640.1translation initiation factor eIF-2B subunit alpha | -0.82 | 4.85e-02 |
| SFRUCORN70000021462 | XP_035433572.2arrestin domain-containing protein 17-like | -0.58 | 4.95e-02 |
| SFRUCORN70000022947 | XP_035453423.1mitochondrial glutamate carrier 1 | -1.30 | 4.95e-02 |

**Table S5.** Wald test_Enriched genes in RC compared to CR

| **GeneID** | **Description** | **Log2FoldChange** | **padj** |
| --- | --- | --- | --- |
| SFRUCORN70000010933 | XP_035446711.1cytoplasmic protein NCK1 isoform X2 | 2.75 | 2.16e-04 |
| SFRUCORN70000025596 | KAG8109051.1hypothetical protein SFRUCORN_015769 | 2.07 | 4.51e-02 |
| SFRUCORN70000019084 | XP_050550216.1mitogen-activated protein kinase kinase kinase 11 isoform X3 | 1.77 | 6.44e-03 |
| SFRUCORN70000027355 | XP_022826470.1MAGUK p55 subfamily member 5 isoform X4 | 1.64 | 4.77e-02 |
| SFRUCORN70000025314 | XP_035456897.1adapter molecule Crk | 2.01 | 2.80e-02 |
| SFRUCORN70000018686 | XP_035431802.2 fat-like cadherin-related tumor suppressor homolog | 1.76 | 4.38e-02 |
| SFRUCORN70000011497 | XP_050557051.1protocadherin-like wing polarity protein stan isoform X1 | 3.11 | 5.80e-06 |
| SFRUCORN70000020674 | XP_035459136.2T-related protein | 1.58 | 4.63e-02 |
| SFRUCORN70000002537 | XP_035452225.11-phosphatidylinositol 4 | 2.17 | 1.43e-02 |
| SFRUCORN70000003408 | XP_035453150.1DE-cadherin isoform X2 | 1.80 | 1.73e-02 |
| SFRUCORN70000018045 | XP_021201251.1hemicentin-2 isoform X2 | 2.13 | 3.63e-02 |
| SFRUCORN70000010605 | XP_035446713.1myosin-VIIa isoform X1 | 2.16 | 1.83e-02 |
| SFRUCORN70000014722 | XP_035430485.1 ichor | 3.36 | 1.26e-08 |
| SFRUCORN70000018215 | XP_035440655.1mothers against decapentaplegic homolog 6 | 1.39 | 8.60e-03 |
| SFRUCORN70000010035 | XP_035435715.1transcription factor sem-2 isoform X2 | 1.93 | 4.96e-02 |
| SFRUCORN70000017239 | KAG8102611.1hypothetical protein SFRUCORN_017369 | 1.65 | 4.39e-02 |
| SFRUCORN70000011943 | XP_022835557.1catenin delta-2 isoform X2 | 1.73 | 3.55e-02 |
| SFRUCORN70000004370 | KAG8116215.1hypothetical protein SFRUCORN_008729 | 1.87 | 2.43e-02 |
| SFRUCORN70000023041 | XP_035429158.2neurotrophin 1 isoform X1 | 1.22 | 1.66e-02 |
| SFRUCORN70000020755 | XP_050560777.1 unconventional myosin-XV | 1.81 | 2.89e-02 |
| SFRUCORN70000004899 | XP_021181254.2spatacsin isoform X1 | 1.91 | 2.96e-03 |
| SFRUCORN70000024638 | XP_035449877.2myosin-I heavy chain isoform X3 | 1.52 | 2.26e-02 |
| SFRUCORN70000020428 | XP_035459151.2kalirin isoform X1 | 1.93 | 1.24e-02 |
| SFRUCORN70000022901 | XP_037302545.1semaphorin-5A isoform X2 | 1.78 | 3.73e-02 |
| SFRUCORN70000010758 | XP_050563932.1tyrosine-protein phosphatase Lar isoform X3 | 2.38 | 3.89e-04 |
| SFRUCORN70000013804 | XP_035445730.1stathmin-4 isoform X3 | 1.20 | 3.73e-02 |
| SFRUCORN70000000723 | XP_035446685.2kelch-like protein 40a | 1.64 | 2.46e-02 |
| SFRUCORN70000030971 | XP_022819216.1neurogenic locus Notch protein isoform X5 | 2.01 | 2.23e-02 |
| SFRUCORN70000014035 | XP_050553614.1exocyst complex component 2 | 1.56 | 2.43e-02 |
| SFRUCORN70000008103 | XP_050562867.1adenomatous polyposis coli protein-like isoform X1 | 1.34 | 4.65e-02 |
| SFRUCORN70000009044 | XP_050563333.1proto-oncogene tyrosine-protein kinase ROS isoform X3 | 2.83 | 8.95e-03 |
| SFRUCORN70000023966 | XP_047036675.1homeobox protein orthopedia-like | 4.16 | 1.05e-06 |
| SFRUCORN70000013710 | XP_049703132.1afadin isoform X2 | 3.29 | 1.28e-06 |
| SFRUCORN70000015678 | XP_035438079.2amyloid beta A4 precursor protein-binding family B member 1-interacting protein isoform X1 | 1.77 | 6.02e-03 |
| SFRUCORN70000005022 | XP_035438907.2leucine-rich melanocyte differentiation-associated protein isoform X2 | 1.99 | 5.17e-04 |
| SFRUCORN70000001970 | XP_050556345.1regulator of G-protein signaling loco isoform X2 | 1.90 | 3.42e-02 |
| SFRUCORN70000022062 | KAG8122324.1hypothetical protein SFRUCORN_013051 | 1.14 | 3.50e-02 |
| SFRUCORN70000012607 | KPI99949.1Homeobox protein abdominal-A-like | 1.86 | 4.47e-02 |
| SFRUCORN70000018214 | XP_035440655.1mothers against decapentaplegic homolog 6 | 1.31 | 4.96e-02 |
| SFRUCORN70000006627 | XP_035433280.1E3 ubiquitin-protein ligase SIAH1 | 1.42 | 1.62e-02 |
| SFRUCORN70000012426 | XP_035447759.2unconventional myosin-IXa isoform X1 | 1.81 | 2.40e-02 |
| SFRUCORN70000020343 | XP_050563731.1unconventional myosin-IXa isoform X2 | 3.26 | 1.08e-03 |
| SFRUCORN70000005173 | XP_035433850.1signal-induced proliferation-associated 1-like protein 1 | 2.10 | 1.13e-02 |
| SFRUCORN70000015996 | XP_035444871.1G1/S-specific cyclin-E | 1.55 | 4.55e-02 |
| SFRUCORN70000033071 | XP_050559313.1spectrin beta chain | 1.74 | 3.31e-02 |
| SFRUCORN70000022620 | XP_035445596.2dedicator of cytokinesis protein 9 isoform X2 | 3.93 | 3.49e-08 |
| SFRUCORN70000009997 | XP_035435862.2CD109 antigen | 1.71 | 1.81e-02 |
| SFRUCORN70000003875 | KAI5633571.1g-protein alpha subunit domain-containing protein | 1.34 | 3.73e-02 |
| SFRUCORN70000003873 | KPJ03305.1Guanine nucleotide-binding protein G(o) subunit alpha 47A | 1.16 | 4.72e-02 |
| SFRUCORN70000029818 | XP_035434904.2CREB-regulated transcription coactivator 3 isoform X1 | 1.53 | 3.81e-02 |
| SFRUCORN70000022092 | KAG8122340.1hypothetical protein SFRUCORN_013067 | 1.54 | 2.90e-03 |
| SFRUCORN70000008325 | KAG8107650.1hypothetical protein SFRUCORN_019853 | 1.59 | 3.84e-02 |
| SFRUCORN70000016933 | XP_050559721.1arf-GAP with coiled-coil | 2.05 | 2.75e-02 |
| SFRUCORN70000004495 | XP_050554467.1active breakpoint cluster region-related protein isoform X1 | 1.76 | 3.40e-02 |
| SFRUCORN70000033877 | XP_035429155.2dedicator of cytokinesis protein 4 isoform X2 | 2.42 | 9.75e-03 |
| SFRUCORN70000015658 | XP_035441798.2ras-specific guanine nucleotide-releasing factor RalGPS2 isoform X3 | 1.67 | 4.26e-02 |
| SFRUCORN70000013382 | XP_035436348.1TBC1 domain family member 13 isoform X3 | 1.80 | 1.72e-02 |
| SFRUCORN70000002508 | XP_035433455.2discoidin domain-containing receptor 2 isoform X2 | 1.67 | 1.93e-02 |
| SFRUCORN70000013913 | XP_032519359.1cytohesin-1 isoform X1 | 2.35 | 1.37e-02 |
| SFRUCORN70000013911 | XP_022831269.1cytohesin-1 isoform X2 | 2.06 | 3.77e-04 |
| SFRUCORN70000004913 | XP_050558958.1Ca(2+)/calmodulin-responsive adenylate cyclase isoform X1 | 1.29 | 4.01e-02 |
| SFRUCORN70000007640 | XP_035437087.1SH3 domain-binding protein 5 homolog isoform X2 | 2.15 | 9.75e-03 |
| SFRUCORN70000015775 | KAI5631504.1cystatin domain-containing protein | 2.49 | 1.23e-02 |
| SFRUCORN70000016644 | XP_035449063.2proteasome activator complex subunit 4B isoform X1 | 1.03 | 4.86e-02 |
| SFRUCORN70000013481 | XP_050562348.1centaurin-gamma-1A isoform X3 | 2.09 | 1.04e-02 |
| SFRUCORN70000027623 | XP_050556305.1serine protease inhibitor A3K | 2.16 | 1.90e-02 |
| SFRUCORN70000021697 | XP_050556279.1DNA replication licensing factor Mcm2 | 1.38 | 3.73e-02 |
| SFRUCORN70000003416 | XP_022824349.1endophilin-A-like isoform X4 | 1.34 | 4.04e-02 |
| SFRUCORN70000010898 | XP_050552074.1transmembrane protein KIAA1109 homolog isoform X6 | 2.57 | 5.80e-03 |
| SFRUCORN70000015867 | XP_035435003.2dnaJ homolog subfamily C member 13 isoform X1 | 1.45 | 4.72e-02 |
| SFRUCORN70000013397 | KAG8114678.1hypothetical protein SFRUCORN_003358 | 1.56 | 1.06e-02 |
| SFRUCORN70000026179 | XP_035444508.1ralBP1-associated Eps domain-containing protein 1 | 1.55 | 1.75e-02 |
| SFRUCORN70000030866 | XP_035444508.1ralBP1-associated Eps domain-containing protein 1 | 1.08 | 4.89e-02 |
| SFRUCORN70000002332 | XP_013193425.1 AP-2 complex subunit mu | 1.27 | 3.39e-02 |
| SFRUCORN70000008653 | XP_035431627.2low-density lipoprotein receptor-related protein 4 | 1.37 | 4.28e-02 |
| SFRUCORN70000000996 | XP_026727826.1flotillin-1 isoform X2 | 1.01 | 4.71e-02 |
| SFRUCORN70000002557 | XP_022832733.1dnaJ homolog subfamily C member 22 isoform X2 | 2.22 | 4.92e-04 |
| SFRUCORN70000019186 | XP_050561044.1epsin-2 isoform X2 | 1.69 | 5.83e-04 |
| SFRUCORN70000020979 | XP_035444778.1low-density lipoprotein receptor-related protein 2 isoform X3 | 1.94 | 1.08e-03 |
| SFRUCORN70000019923 | XP_050556783.1endoribonuclease Dcr-1 isoform X1 | 2.07 | 2.75e-02 |
| SFRUCORN70000004775 | KAG8117383.1hypothetical protein SFRUCORN_019532 | 1.53 | 2.00e-02 |
| SFRUCORN70000009019 | XP_035450081.2endoribonuclease Dicer | 2.47 | 2.96e-03 |
| SFRUCORN70000006571 | XP_035432811.1protein Gawky isoform X1 | 2.24 | 1.14e-02 |
| SFRUCORN70000006159 | XP_050551215.1putative helicase mov-10-B.1 | 2.94 | 3.45e-03 |

**Table S6.** Wald test_Enriched genes in CR compared to RC

| **GeneID** | **Description** | **Log2FoldChange** | **padj** |
| --- | --- | --- | --- |
| SFRUCORN70000014653 | XP_035447279.239S ribosomal protein L34 | -1.25 | 0.02 |
| SFRUCORN70000015711 | XP_035446957.139S ribosomal protein L2 | -1.08 | 0.03 |
| SFRUCORN70000024005 | XP_035433513.128S ribosomal protein S21 | -1.19 | 0.02 |
| SFRUCORN70000017995 | XP_035433985.139S ribosomal protein L4 | -1.12 | 0.03 |
| SFRUCORN70000026545 | XP_050559619.1probable proline--tRNA ligase | -1.17 | 0.04 |
| SFRUCORN70000014287 | XP_050559749.139S ribosomal protein L43 | -1.53 | 0.01 |
| SFRUCORN70000029995 | XP_035457282.228S ribosomal protein S35 | -1.13 | 0.04 |
| SFRUCORN70000024903 | XP_035446460.1la protein homolog | -1.22 | 0.01 |
| SFRUCORN70000000508 | XP_035446371.128S ribosomal protein S18b | -1.41 | 0.01 |
| SFRUCORN70000000736 | XP_035446053.139S ribosomal protein L15 | -1.21 | 0.02 |
| SFRUCORN70000010722 | XP_035445897.139S ribosomal protein L3 | -1.02 | 0.05 |
| SFRUCORN70000010621 | XP_035446446.1elongation factor Ts | -1.04 | 0.03 |
| SFRUCORN70000003502 | XP_035430007.139S ribosomal protein L51 | -1.17 | 0.04 |
| SFRUCORN70000006456 | XP_026726176.1elongation factor 1-alpha 2 | -1.47 | 0.03 |
| SFRUCORN70000024289 | XP_035429888.228S ribosomal protein S14 | -1.34 | 0.02 |
| SFRUCORN70000024290 | XP_022828292.160S acidic ribosomal protein P1 | -1.26 | 0.04 |
| SFRUCORN70000025533 | XP_022824544.160S acidic ribosomal protein P0 | -1.21 | 0.04 |
| SFRUCORN70000033907 | ASN63931.1glutathione S-transferase delta 2 | -1.74 | 0.02 |
| SFRUCORN70000006498 | XP_035440670.228S ribosomal protein S24 | -1.21 | 0.05 |
| SFRUCORN70000017282 | XP_035440640.1translation initiation factor eIF-2B subunit alpha | -1.04 | 0.04 |
| SFRUCORN70000027369 | CAH2234230.1 jg16679 | -1.39 | 0.02 |
| SFRUCORN70000033432 | CAH2234230.1 jg16679 | -1.41 | 0.01 |
| SFRUCORN70000003922 | XP_035434673.228S ribosomal protein S17 | -1.24 | 0.05 |
| SFRUCORN70000003791 | XP_035435176.228S ribosomal protein S9 | -1.25 | 0.01 |
| SFRUCORN70000026212 | XP_022823274.140S ribosomal protein S26 | -1.26 | 0.04 |
| SFRUCORN70000026220 | XP_021195351.160S ribosomal protein L34 | -1.26 | 0.04 |
| SFRUCORN70000004653 | XP_035439403.228S ribosomal protein S18a | -1.20 | 0.03 |
| SFRUCORN70000021850 | XP_022827032.140S ribosomal protein S12 | -1.26 | 0.03 |
| SFRUCORN70000019080 | XP_035437304.139S ribosomal protein L33 | -1.37 | 0.03 |
| SFRUCORN70000027077 | XP_035437342.239S ribosomal protein L17 | -1.11 | 0.04 |
| SFRUCORN70000022189 | XP_035437284.2ribosome-recycling factor | -1.14 | 0.03 |
| SFRUCORN70000022210 | XP_035437439.139S ribosomal protein L14 | -1.45 | 0.01 |
| SFRUCORN70000008960 | XP_022831736.160S acidic ribosomal protein P2 | -1.26 | 0.05 |
| SFRUCORN70000011863 | XP_035448020.128S ribosomal protein S15 | -1.26 | 0.03 |
| SFRUCORN70000008703 | XP_050563190.1FAU ubiquitin-like and ribosomal protein S30 | -1.25 | 0.03 |
| SFRUCORN70000006870 | XP_021187435.140S ribosomal protein S16 isoform X2 | -1.27 | 0.03 |
| SFRUCORN70000020520 | XP_035442579.160S ribosomal protein L28 | -1.23 | 0.04 |
| SFRUCORN70000002071 | XP_035454757.2probable 39S ribosomal protein L24 | -1.28 | 0.01 |
| SFRUCORN70000001379 | XP_035454812.2probable 28S ribosomal protein S16 | -1.23 | 0.03 |
| SFRUCORN70000025669 | XP_035454966.2elongation factor Tu-like | -1.06 | 0.05 |
| SFRUCORN70000009698 | KOB63048.140S ribosomal protein S11 | -1.13 | 0.05 |
| SFRUCORN70000030339 | XP_035458222.160S ribosomal protein L27a | -1.19 | 0.04 |
| SFRUCORN70000008833 | KAI5634320.1ribosomal l39 protein domain-containing protein | -1.32 | 0.02 |
| SFRUCORN70000029866 | XP_035431121.239S ribosomal protein L22 | -1.04 | 0.04 |
| SFRUCORN70000024457 | XP_035429056.139S ribosomal protein L20 | -2.27 | 0.00 |
| SFRUCORN70000010104 | XP_021190682.140S ribosomal protein S24 | -1.19 | 0.05 |
| SFRUCORN70000020726 | AAK72378.160S ribosomal protein L7/L12 precursor | -1.16 | 0.05 |
| SFRUCORN70000032674 | XP_035432546.128S ribosomal protein S7 | -1.12 | 0.04 |
| SFRUCORN70000004158 | XP_035432950.2translation factor GUF1 homolog | -1.12 | 0.04 |
| SFRUCORN70000004269 | XP_050560970.140S ribosomal protein SA | -1.25 | 0.04 |
| SFRUCORN70000006628 | XP_021183306.160S ribosomal protein L29 | -1.25 | 0.03 |
| SFRUCORN70000031076 | XP_035458385.2protein penguin | -1.39 | 0.01 |
| SFRUCORN70000019191 | XP_035433187.2peptidyl-tRNA hydrolase ICT1 | -1.16 | 0.04 |
| SFRUCORN70000004011 | XP_035436624.239S ribosomal protein L21 | -1.22 | 0.03 |
| SFRUCORN70000022839 | XP_035436416.139S ribosomal protein L16 | -1.30 | 0.02 |
| SFRUCORN70000004981 | XP_035436173.1elongation factor Tu | -1.03 | 0.02 |
| SFRUCORN70000019593 | XP_035449887.139S ribosomal protein L11 | -1.30 | 0.02 |
| SFRUCORN70000005572 | XP_035449920.1transmembrane emp24 domain-containing protein 5 | -1.29 | 0.02 |
| SFRUCORN70000005515 | XP_035449524.128S ribosomal protein S5 | -1.00 | 0.02 |
| SFRUCORN70000012698 | XP_035441762.139S ribosomal protein L35 | -1.21 | 0.02 |
| SFRUCORN70000018383 | XP_035442109.228S ribosomal protein S18c | -1.57 | 0.01 |
| SFRUCORN70000030184 | XP_050562451.1peptide deformylase | -1.63 | 0.02 |
| SFRUCORN70000002129 | XP_035438333.139S ribosomal protein L32 | -1.11 | 0.04 |
| SFRUCORN70000020034 | XP_022815642.1DPH3 homolog isoform X1 | -1.48 | 0.01 |
| SFRUCORN70000024710 | XP_035452842.139S ribosomal protein L19 | -1.09 | 0.04 |
| SFRUCORN70000007512 | OWR54112.140S ribosomal protein S28 | -1.18 | 0.05 |
| SFRUCORN70000007515 | XP_035445032.2alanine--glyoxylate aminotransferase 2 | -1.37 | 0.00 |
| SFRUCORN70000007533 | XP_050549496.139S ribosomal protein L52 | -1.20 | 0.04 |
| SFRUCORN70000027235 | XP_026499221.154S ribosomal protein c83.06c | -1.31 | 0.01 |
| SFRUCORN70000025166 | XP_013171955.1 28S ribosomal protein S34 | -1.22 | 0.01 |
| SFRUCORN70000009924 | XP_035454829.1probable dimethyladenosine transferase | -1.35 | 0.02 |
| SFRUCORN70000013878 | XP_035446298.1nucleolar protein 58 | -1.16 | 0.02 |
| SFRUCORN70000000587 | XP_050549685.1nucleolar protein 10 isoform X6 | -1.00 | 0.04 |
| SFRUCORN70000020218 | XP_035430923.1periodic tryptophan protein 1 homolog | -1.34 | 0.01 |
| SFRUCORN70000008073 | XP_035440725.2protein Peter pan | -1.19 | 0.03 |
| SFRUCORN70000018637 | XP_035435752.2protein KRI1 homolog | -1.32 | 0.02 |
| SFRUCORN70000013262 | XP_035452286.2nucleolar protein 9 | -1.25 | 0.02 |
| SFRUCORN70000016941 | XP_035439538.2ESF1 homolog | -0.97 | 0.04 |
| SFRUCORN70000004500 | XP_035439934.2notchless protein homolog 1 | -0.91 | 0.04 |
| SFRUCORN70000005938 | XP_050560788.1probable RNA 3'-terminal phosphate cyclase-like protein | -1.12 | 0.05 |
| SFRUCORN70000005960 | XP_035439867.1complement component 1 Q subcomponent-binding protein | -1.56 | 0.00 |
| SFRUCORN70000031308 | XP_050550174.1ribosome biogenesis protein BOP1 homolog | -1.37 | 0.03 |
| SFRUCORN70000020261 | XP_050552736.1 39S ribosomal protein L44 | -1.23 | 0.04 |
| SFRUCORN70000017690 | XP_022828307.1ribosome production factor 2 homolog | -1.10 | 0.04 |
| SFRUCORN70000008684 | XP_050563260.1nucleolar protein 16 | -1.04 | 0.04 |
| SFRUCORN70000008982 | XP_022832149.14-coumarate--CoA ligase-like 5 | -1.03 | 0.03 |
| SFRUCORN70000024160 | XP_004933552.1U6 snRNA-associated Sm-like protein LSm6 | -1.14 | 0.05 |
| SFRUCORN70000008807 | XP_035449567.2H/ACA ribonucleoprotein complex subunit 3 | -1.20 | 0.04 |
| SFRUCORN70000026498 | XP_050561467.1protein MAK16 homolog B | -1.42 | 0.01 |
| SFRUCORN70000021271 | XP_050561522.1periodic tryptophan protein 2 homolog | -1.03 | 0.03 |
| SFRUCORN70000023582 | XP_035451304.1rRNA 2'-O-methyltransferase fibrillarin | -1.22 | 0.02 |
| SFRUCORN70000030416 | XP_050557301.1pre-rRNA-processing protein TSR1 homolog | -0.95 | 0.03 |
| SFRUCORN70000023908 | XP_035429088.1probable rRNA-processing protein EBP2 homolog | -1.30 | 0.03 |
| SFRUCORN70000032670 | XP_035432798.2transducin beta-like protein 3 | -1.03 | 0.05 |
| SFRUCORN70000007250 | XP_022819847.1pre-rRNA processing protein FTSJ3 | -1.09 | 0.04 |
| SFRUCORN70000004872 | XP_022819756.125S rRNA (cytosine-C(5))-methyltransferase nop2 | -1.04 | 0.04 |
| SFRUCORN70000005085 | XP_035438584.2MKI67 FHA domain-interacting nucleolar phosphoprotein | -0.93 | 0.04 |
| SFRUCORN70000019996 | XP_035453169.1ribosome biogenesis protein NOP53 | -1.16 | 0.02 |
| SFRUCORN70000003353 | XP_035452696.2PIN2/TERF1-interacting telomerase inhibitor 1 | -0.97 | 0.04 |
| SFRUCORN70000020958 | XP_022829790.1NHP2-like protein 1 | -1.39 | 0.03 |
| SFRUCORN70000015148 | XP_035444971.2U3 small nucleolar RNA-associated protein 6 homolog | -1.03 | 0.05 |
| SFRUCORN70000020148 | XP_035444632.2dimethyladenosine transferase 1 | -1.15 | 0.02 |
| SFRUCORN70000022552 | XP_035434502.2cytochrome c oxidase subunit 7A2 | -1.33 | 0.02 |
| SFRUCORN70000005188 | XP_022815137.1ATP synthase subunit delta | -1.15 | 0.05 |
| SFRUCORN70000010586 | XP_035452545.1cytochrome c oxidase subunit 5A | -1.17 | 0.03 |
| SFRUCORN70000007745 | XP_035454040.2cytochrome b-c1 complex subunit 7 | -1.14 | 0.05 |
| SFRUCORN70000016418 | XP_035453364.1ATP synthase subunit g | -1.37 | 0.02 |
| SFRUCORN70000017149 | XP_035453819.1ATP synthase subunit e | -1.24 | 0.04 |
| SFRUCORN70000029744 | XP_035456104.1NADH dehydrogenase [ubiquinone] iron-sulfur protein 2 | -0.95 | 0.03 |
| SFRUCORN70000000553 | XP_035446663.1probable NADH dehydrogenase [ubiquinone] iron-sulfur protein 6 | -1.12 | 0.04 |
| SFRUCORN70000027331 | XP_035435039.1ATP synthase subunit alpha | -1.15 | 0.02 |
| SFRUCORN70000005270 | XP_021188571.2serine protease nudel | -1.27 | 0.02 |
| SFRUCORN70000028785 | XP_022818617.1cytochrome c oxidase subunit 5B | -1.38 | 0.01 |
| SFRUCORN70000004534 | XP_035440005.1cytochrome c oxidase subunit 6A | -1.21 | 0.03 |
| SFRUCORN70000011228 | XP_035447994.2NADH dehydrogenase [ubiquinone] 1 alpha subcomplex subunit 7-like | -1.06 | 0.05 |
| SFRUCORN70000017718 | XP_022828266.1ATP synthase subunit O | -1.15 | 0.04 |
| SFRUCORN70000029864 | XP_035431122.1cytochrome b-c1 complex subunit 8 | -1.17 | 0.04 |
| SFRUCORN70000031945 | XP_035459089.1protein stunted isoform X1 | -1.18 | 0.04 |
| SFRUCORN70000010108 | XP_035432545.2NADH-ubiquinone oxidoreductase subunit 8 | -1.16 | 0.03 |
| SFRUCORN70000033667 | KAG7158486.1NADH dehydrogenase subunit 3-like | -1.76 | 0.01 |
| SFRUCORN70000033671 | XP_037877556.1 cytochrome c oxidase subunit 1-like | -1.34 | 0.01 |
| SFRUCORN70000032706 | XP_035449824.1ATP synthase-coupling factor 6 | -1.30 | 0.02 |
| SFRUCORN70000002161 | XP_035438428.1ATP synthase subunit beta | -1.24 | 0.01 |
| SFRUCORN70000002165 | XP_035438473.1ATP synthase subunit beta | -1.17 | 0.02 |
| SFRUCORN70000007477 | XP_035444883.1protein trapped in endoderm-1 isoform X2 | -1.08 | 0.05 |
| SFRUCORN70000017998 | XP_035433956.1NADH dehydrogenase [ubiquinone] 1 subunit C2 | -1.24 | 0.02 |
| SFRUCORN70000009923 | XP_050682055.1NADH dehydrogenase [ubiquinone] 1 beta subcomplex subunit 6 | -1.14 | 0.04 |
| SFRUCORN70000003646 | XP_022828183.160S ribosomal protein L30 | -1.15 | 0.05 |
| SFRUCORN70000028336 | XP_050562941.150S ribosomal protein L1 | -1.38 | 0.02 |
| SFRUCORN70000027136 | XP_035441904.228S ribosomal protein S10 | -1.39 | 0.03 |
| SFRUCORN70000009920 | XP_035445623.1NADH dehydrogenase [ubiquinone] 1 alpha subcomplex subunit 5 | -1.11 | 0.04 |
| SFRUCORN70000021040 | XP_035454456.1V-type proton ATPase subunit e 2 | -1.30 | 0.03 |
| SFRUCORN70000017554 | XP_035458777.1cytochrome b-c1 complex subunit 10 | -1.23 | 0.03 |
| SFRUCORN70000014174 | XP_026756379.1cytochrome c oxidase subunit 7C | -1.28 | 0.02 |
| SFRUCORN70000004188 | XP_035432112.2NADH dehydrogenase [ubiquinone] 1 beta subcomplex subunit 10 | -1.05 | 0.05 |
| SFRUCORN70000020464 | UAJ21579.1 V-ATPase subunit B | -0.86 | 0.04 |
| SFRUCORN70000005020 | XP_035438547.2NADH dehydrogenase [ubiquinone] 1 beta subcomplex subunit 2 | -1.25 | 0.02 |
| SFRUCORN70000029355 | XP_035444807.1cytochrome b-c1 complex subunit 2 | -1.03 | 0.04 |
| SFRUCORN70000011419 | XP_035444327.1adenylate kinase isoenzyme 6 homolog | -1.38 | 0.01 |
| SFRUCORN70000005949 | XP_022814095.1acyl carrier protein | -1.18 | 0.03 |
| SFRUCORN70000000066 | XP_035444917.2ribonucleases P/MRP protein subunit POP1 | -1.10 | 0.04 |
| SFRUCORN70000016322 | XP_035457750.2guanine nucleotide-binding protein-like 3 homolog | -1.02 | 0.03 |
